# SLIM: A small linear model with STRING embeddings for single-cell genetic perturbation prediction

**DOI:** 10.64898/2026.08.07.743481

**Authors:** Dewei Hu, Marc Pielies Avellí, Lars Juhl Jensen, Simon Rasmussen

## Abstract

Predicting cellular responses to genetic perturbations is central to understanding gene function and prioritizing therapeutic targets, but experimental screens cannot exhaustively cover genes, cell types, and perturbation combinations. Recent benchmarks have shown that simple baselines can match or outperform substantially more complex models, suggesting that informative biological priors may be as important as model capacity. Here we present SLIM, a lightweight extension of the bilinear model of Ahlmann-Eltze et al. SLIM represents perturbations with 64-dimensional embeddings derived from the STRING protein network and predicts mean transcriptional responses through a closed-form ridge-regression estimator. It then constructs single-cell populations by retrieving training cells and rescaling each gene to match the predicted mean. We evaluated SLIM against four deep learning models and two simple baselines on four single-gene perturbation datasets and one combinatorial perturbation dataset. Across these within-dataset benchmarks, SLIM achieved competitive mean-response accuracy, ranked first in eight of twelve single-gene dataset–metric comparisons, and produced substantially lower maximum mean discrepancy values than the evaluated alternatives. The model has 640 trainable parameters and fitted each benchmark dataset in under 10 seconds on a CPU. These results show that compact biological representations can support accurate and computationally efficient perturbation prediction. Code is available at https://github.com/RasmussenLab/SLIM.

**Key Points:**

- SLIM combines a closed-form bilinear predictor with STRING-derived perturbation embeddings.
- Across five within-dataset benchmarks, SLIM achieved competitive mean-response prediction with only 640 trainable parameters.
- SLIM builds cell populations by rescaling retrieved training cells to the predicted mean, so they inherit realistic cell-to-cell variation and gene–gene covariation.
- The results highlight the importance of perturbation representations and population-construction procedures in low-data benchmarks.
- SLIM fits each benchmark dataset in under 10 seconds on a standard CPU.

## Introduction

Understanding how individual cells respond to genetic perturbations is central to deciphering gene function, dissecting the molecular basis of disease, and identifying therapeutic targets [1]. High-throughput single-cell perturbation screens, such as Perturb-seq [2], now make it possible to measure transcriptome-wide responses to hundreds of genetic knockouts in a single pooled screen. Despite this scale, exhaustive coverage of the perturbation space across all genes, cell types, and disease contexts remains prohibitively expensive. Therefore, computational models capable of predicting cellular responses to unmeasured genes and unseen conditions based on a limited set of observed perturbations are highly desirable [3].

A growing number of deep learning models have been developed for this task. These models employ either data-driven representations from large-scale single-cell datasets or external biological knowledge, such as interaction networks and textual descriptions of gene functions. CPA [4] is a conditional variational autoencoder that encodes perturbations as learned categorical embeddings and models their effects as additive shifts in a disentangled latent space, enabling composition at inference time to predict unseen combinations. GEARS [5] propagates gene embeddings through a graph neural network built over two biological knowledge graphs (gene co-expression and Gene Ontology), capturing non-additive interaction effects for combinatorial knockouts. scGPT [6] is a transformer foundation model pre-trained on millions of single-cell profiles and can be fine-tuned for perturbation prediction. scLAMBDA [7] is a disentangled variational autoencoder that represents perturbed genes through GenePT embeddings [8], which are large language model encodings of textual gene function descriptions. These models rely on expressive architectures and rich biological priors to capture complex perturbation effects.

Recent systematic benchmarks have challenged the assumption that greater architectural complexity necessarily improves perturbation prediction. Ahlmann-Eltze et al. [9] showed that a bilinear model based on perturbation-context PCA representations can match or outperform deep learning approaches across several datasets and metrics, and even the training-set mean was a competitive baseline. In a broader evaluation, Wei et al. [10] found that foundation models often underperform simpler baselines when fine-tuning data are limited and provide gains only when sufficiently large training sets are available. Models incorporating prior knowledge also consistently outperformed those relying on transcriptome-derived embeddings alone, identifying the perturbation representation as an important determinant of generalisation. Other recent studies likewise support the value of prior knowledge for single-cell perturbation prediction [11, 12, 13].

Together, these findings motivate a simple model paired with a biologically informed perturbation representation. One such source of prior knowledge is the STRING database. STRING scores physical interactions and functional associations between proteins, integrating genomic context, high-throughput experiments, conserved co-expression, text mining, and curated databases [14]. Much of this evidence comes from curated databases and text mining and is not present in expression data at all, so models pretrained on expression cannot learn it regardless of how many cells they see. Embeddings of the STRING network are therefore a natural choice of perturbation representation [15, 16]. They have supported protein function prediction, pathway prediction, and disease protein prediction, so we reasoned they carry transferable information about perturbations not observed during training.

Here we present SLIM, which extends the bilinear framework of Ahlmann-Eltze et al. [9] by using STRING embeddings as perturbation-specific priors. SLIM separates mean-response prediction from population reconstruction. A bilinear map between a low-dimensional, context-specific PCA basis and 64-dimensional STRING embeddings predicts the mean transcriptional response, with model parameters estimated in closed form. The model predicts a mean expression profile for each perturbation. To recover cell-level variation, we sample real training cells and scale their gene-wise means to the predicted values, which preserves the observed spread while relocating the distribution. We benchmarked SLIM on single-gene perturbation from K562, RPE1, HepG2, and Jurkat datasets [2, 17] and on the Norman et al. combinatorial overexpression dataset [18]. Across these within-dataset evaluations, SLIM was competitive with the strongest deep learning model for mean-response metrics and achieved the lowest MMD values, while fitting each dataset in under 10 seconds on a CPU.

## Results

### Small linear model with STRING embeddings and retrieval-augmented sampling

SLIM extends the bilinear framework of Ahlmann-Eltze et al. to predict pseudobulk expression profiles for genetic perturbations (**Y**_train_). We first apply PCA to baseline-centred expression profiles across training perturbations (Figure 1). The resulting matrix **G** is a low-dimensional, dataset-specific representation of how readout genes co-vary in the cellular context. We represent each perturbed gene by a node2vec embedding derived from STRING v12.0 [14, 19], as described previously [16]. These protein-level embeddings form **P** and encode functional relationships that can be used for perturbations not observed during training. A single trainable matrix **W** maps between the STRING perturbation space and the fixed PCA readout space, while ***b*** represents the perturbation-independent mean expression profile.

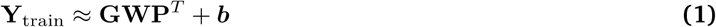

**Figure 1.**
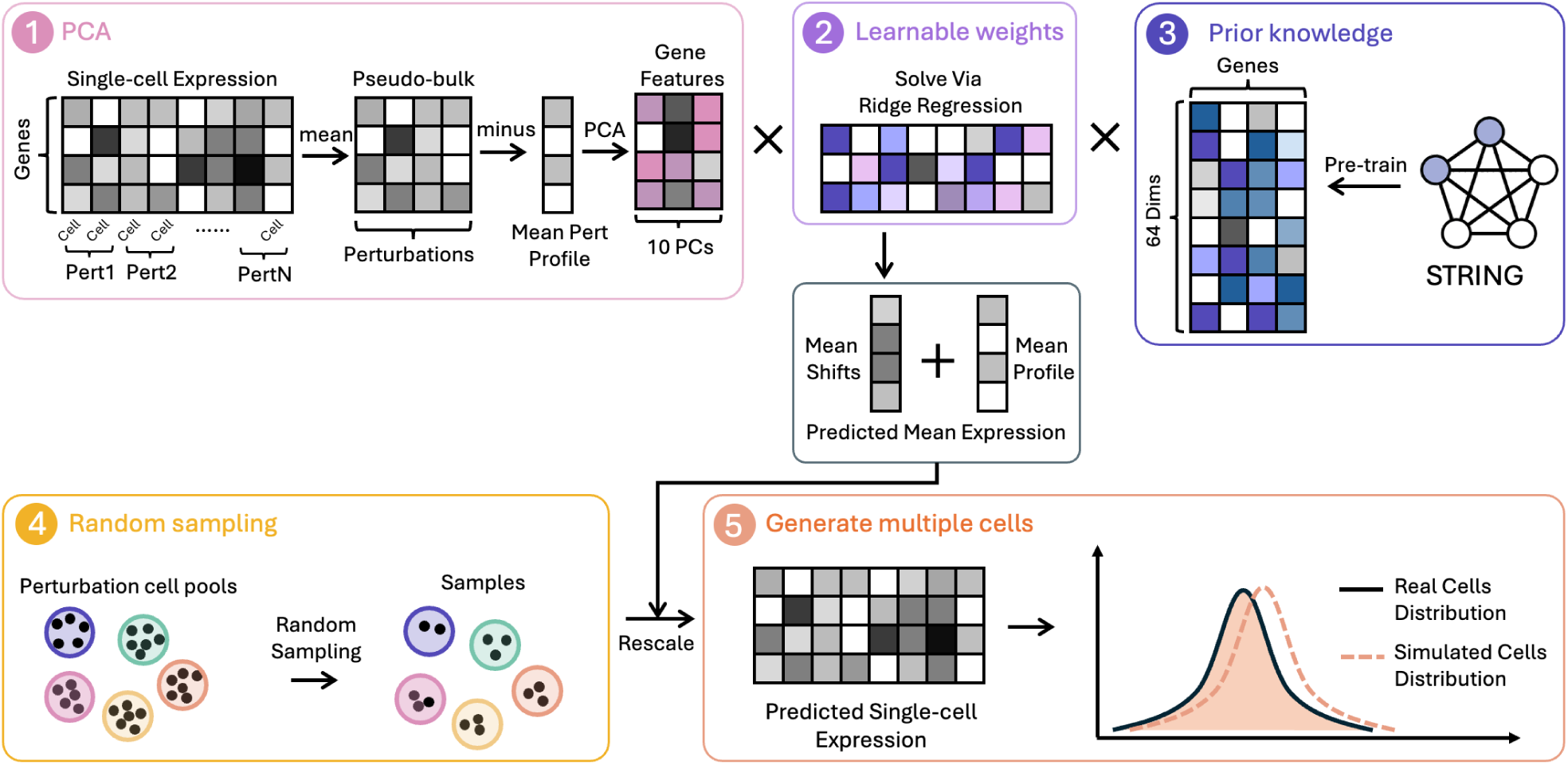
Overview of SLIM. The SLIM pipeline consists of five steps. **(1) PCA:** Single-cell expression profiles (*∼*20k genes *×* cells) are mean-centered and decomposed via PCA to yield 10 principal components per gene, providing a compact, context-aware readout of gene representation. **(2) Learnable weights:** A bilinear weight matrix maps the PCA gene space to the STRING embedding space; the resulting mean shifts are added to the mean expression profile to produce the predicted mean expression for each perturbation. **(3) Prior knowledge:** Each perturbed gene is represented by a 64-dimensional embedding pre-trained on the STRING protein–protein interaction network, supplying biology-informed priors for unseen perturbations. **(4) Random sampling:** Cells are drawn at random from the perturbation cell pools in the training data. **(5) Generate multiple cells:** The sampled cells are multiplicatively rescaled gene-by-gene so that each gene’s mean across the synthetic population matches the predicted mean for that gene, yielding a single-cell population whose distribution closely matches that of real perturbed cells.

The model is fitted using a closed-form ridge-regression estimator, avoiding iterative gradient-based optimisation.

To obtain cell-level outputs from a model trained on perturbation means, we introduce a retrieval-augmented rescaling strategy. For each test perturbation, cells are sampled from the training pool and multiplicatively rescaled so that their gene-wise population means match the SLIM prediction. This procedure preserves heterogeneity and gene–gene covariation present in the retrieved cells, transferring this variation from the training population rather than generating perturbation-specific variability de novo.

### SLIM is competitive on mean-response metrics and yields low MMD in single-gene perturbation prediction

Single-gene perturbation prediction is the core benchmark for evaluating how well a model generalises to unseen knockouts. Given the limited number of training perturbations available in any single dataset, we asked whether a simple linear model coupled with biology-informed representations could match or exceed the performance of substantially larger deep learning models across diverse cell types and experimental conditions.

We benchmarked SLIM against scLAMBDA, GEARS, scGPT, CPA, and the linear model from Ahlmann-Eltze et al. [9] on four single-gene datasets from K562, RPE1, HepG2, and Jurkat cells using Pearson Delta, MMD, and MAE (Figure 2). To study the benefit of our random sampling and rescaling strategy, we added SLIM-Gaussian. SLIM-Gaussian uses the same predicted means as SLIM, but it expands them by sampling each gene independently from a Gaussian with the predicted mean and the training-set per-gene standard deviation. We used the same Gaussian sampling for other mean-only methods (linear model and GEARS).

**Figure 2.**
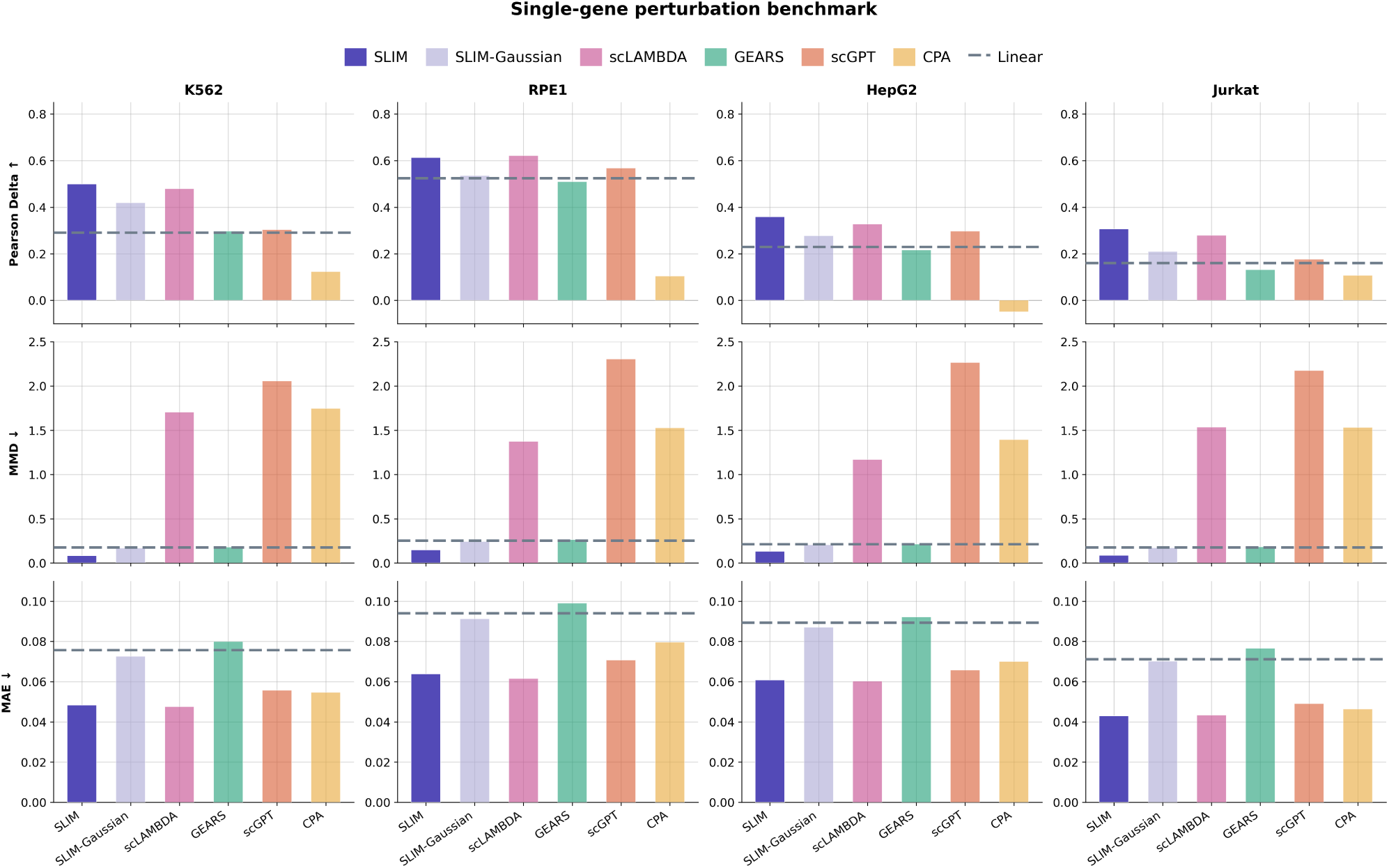
Single-gene perturbation benchmark across four cell lines. Per-dataset performance of SLIM and competing methods on Pearson Delta (higher is better), MMD (lower is better), and MAE (lower is better) across K562, RPE1, HepG2, and Jurkat datasets. Dashed lines indicate the linear baselines.

SLIM ranked first in eight of the twelve dataset–metric comparisons and second in the remaining four, including three in MAE and one in Pearson Delta on RPE1 (Figure S1). Its mean-response performance was similar to that of scLAMBDA: Pearson Delta ranged from 0.31 to 0.61 for SLIM and from 0.28 to 0.62 for scLAMBDA, while MAE ranged from 0.043 to 0.064 and from 0.043 to 0.062, respectively. The largest numerical differences occurred for MMD. SLIM obtained values of 0.080–0.15, compared with 1.2–1.7 for scLAMBDA, 0.18–0.26 for GEARS, and 2.1–2.3 for scGPT. This result is consistent with the design of the retrieval procedure, which preserves multivariate structure from observed training cells. Because the linear baselines instead used independent per-gene Gaussian sampling, the MMD comparison reflects both mean-response accuracy and the population-construction strategy. As a comparison, SLIM-Gaussian shows generally worse scores than SLIM across the datasets and metrics. The per-perturbation distributions underlying these averages are shown in Figure S8.

Together, these results show that a compact linear model with good embedding priors can match substantially larger models in the evaluated low-data, within-context setting. They also show that the procedure used to construct cell populations strongly affects distributional metrics such as MMD.

### Performance varies across combinatorial perturbation settings

Double-gene perturbation prediction is a substantially harder problem than single-gene prediction. The number of gene pairs grows quadratically with the number of genes, and pairs can produce non-additive interaction effects that are difficult to predict from individual knockouts. To test whether SLIM generalised to combinatorial perturbations, we benchmarked it on the Norman dataset [18], which contains overexpression (CRISPRa) combinations in K562 cells. The dataset defines four splits of increasing difficulty based on how many of the two perturbed genes were seen as single perturbations during training: combo seen 2 (both genes seen individually), combo seen 1 (one seen), combo seen 0 (neither seen), and unseen single (a single-gene perturbation held out entirely). To adapt our model for double perturbations, we averaged the embeddings of the two given genes, while we kept the original embeddings for single perturbations. We also used different biases, *b*_single_ and *b*_double_, for single and double perturbations.

Across the four splits and three metrics, SLIM had a median rank of one, although its performance varied more than in the single-gene benchmarks (Figure S2). On the double-perturbation splits, SLIM achieved lower MAE than the competing methods (0.015–0.020, compared with 0.020–0.023 for scLAMBDA) and matched or exceeded scLAMBDA on Pearson Delta in two of the three splits (0.67–0.77 versus 0.60–0.79). scLAMBDA performed better on the combo-seen-0 split. The unseen-single split was more challenging: SLIM achieved a Pearson Delta of 0.38, below all the other benchmarked models, and its MAE also deteriorated relative to the strongest models. Thus, SLIM performed well when predicting combinations but was less effective when extrapolating to an entirely unseen single-gene perturbation in this mixed single- and double-perturbation setting. MMD in all splits remained lower for SLIM (0.10–0.20) than for all the other models (0.31–2.2); Figure 3), consistent with the effect of retrieval-based population construction. As a comparison, SLIM-Gaussian shows generally worse scores than SLIM across the splits and metrics, but is a robust model. The corresponding per-perturbation distributions are shown in Figure S9.

**Figure 3.**
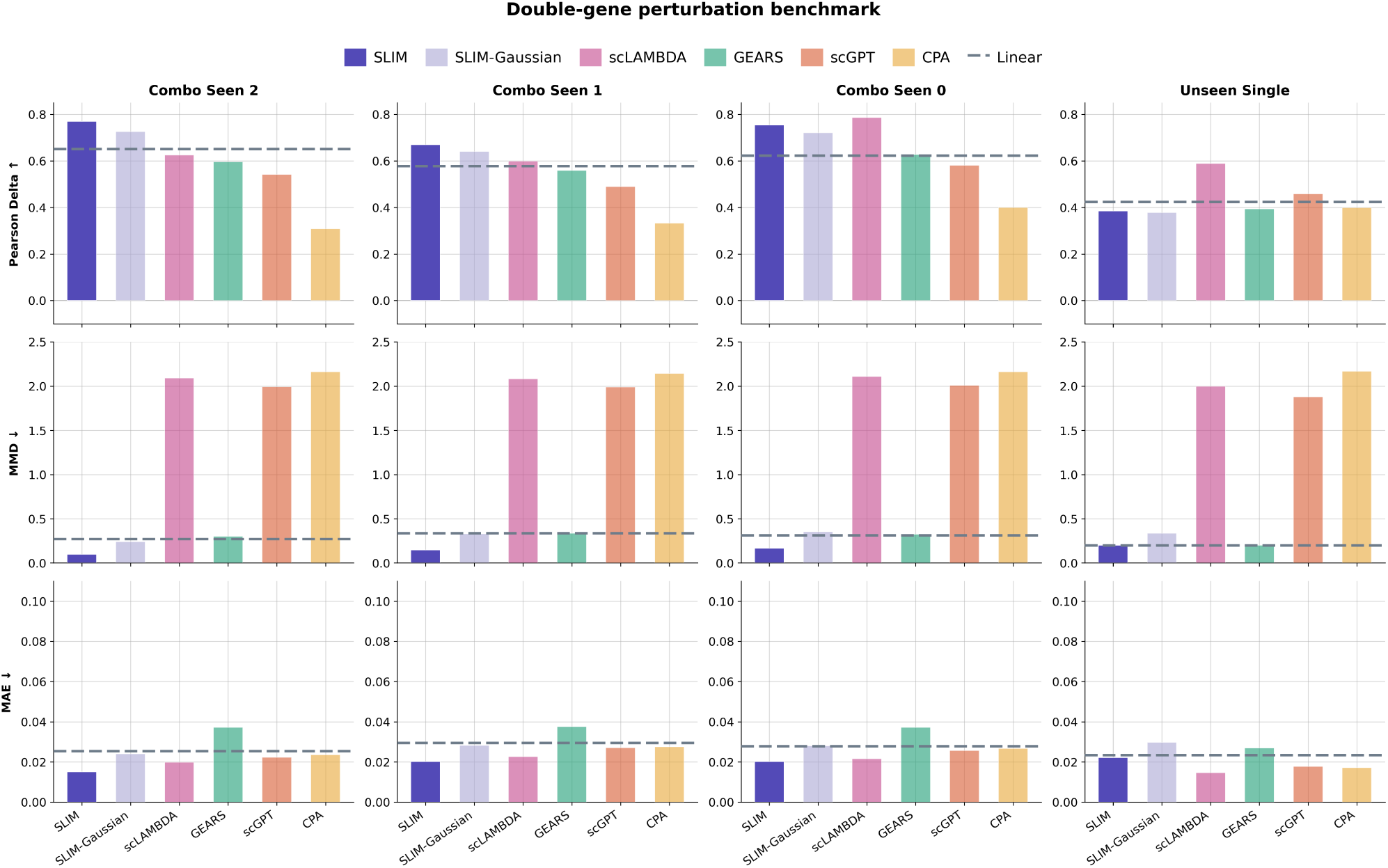
Double-gene perturbation benchmark on the Norman dataset. Per-split performance on Pearson Delta (higher is better), MMD (lower is better), and MAE (lower is better) across Combo Seen 2, 1, and 0, indicating how many constituent genes were observed during training, and Unseen Single, comprising held-out single-gene perturbations. Dashed lines indicate the linear baselines.

These results show that the same model works well in the combinatorial setting without architectural changes, while extrapolation to an entirely unseen single gene remains its weakest case. The random sampling and rescaling method also improved the MMD by keeping the real cell population structure.

### Ablation of hyperparameters and biological priors

SLIM has two hyperparameters: the number of principal components *K* in the compressed readout representation **G** and the ridge regularisation strength *λ*. The STRING embedding dimension, *D* = 64, is fixed. We selected *K* = 10 and *λ* = 0.1 using the validation sets (Figures S6 and S7) and used this configuration for all primary test-set comparisons. We subsequently performed a post hoc sensitivity analysis on the held-out test sets using *K* ∈ {5, 10, 20, 50} and *λ* ∈ {0.01, 0.1, 1, 10} (Figures 4 and 5). Across the four single-gene datasets, all three metrics changed little over the evaluated ranges, and the prespecified configuration lay within this broad region of stable performance. Because these sweeps used test data, they were used only to assess sensitivity and did not influence model selection.

**Figure 4.**
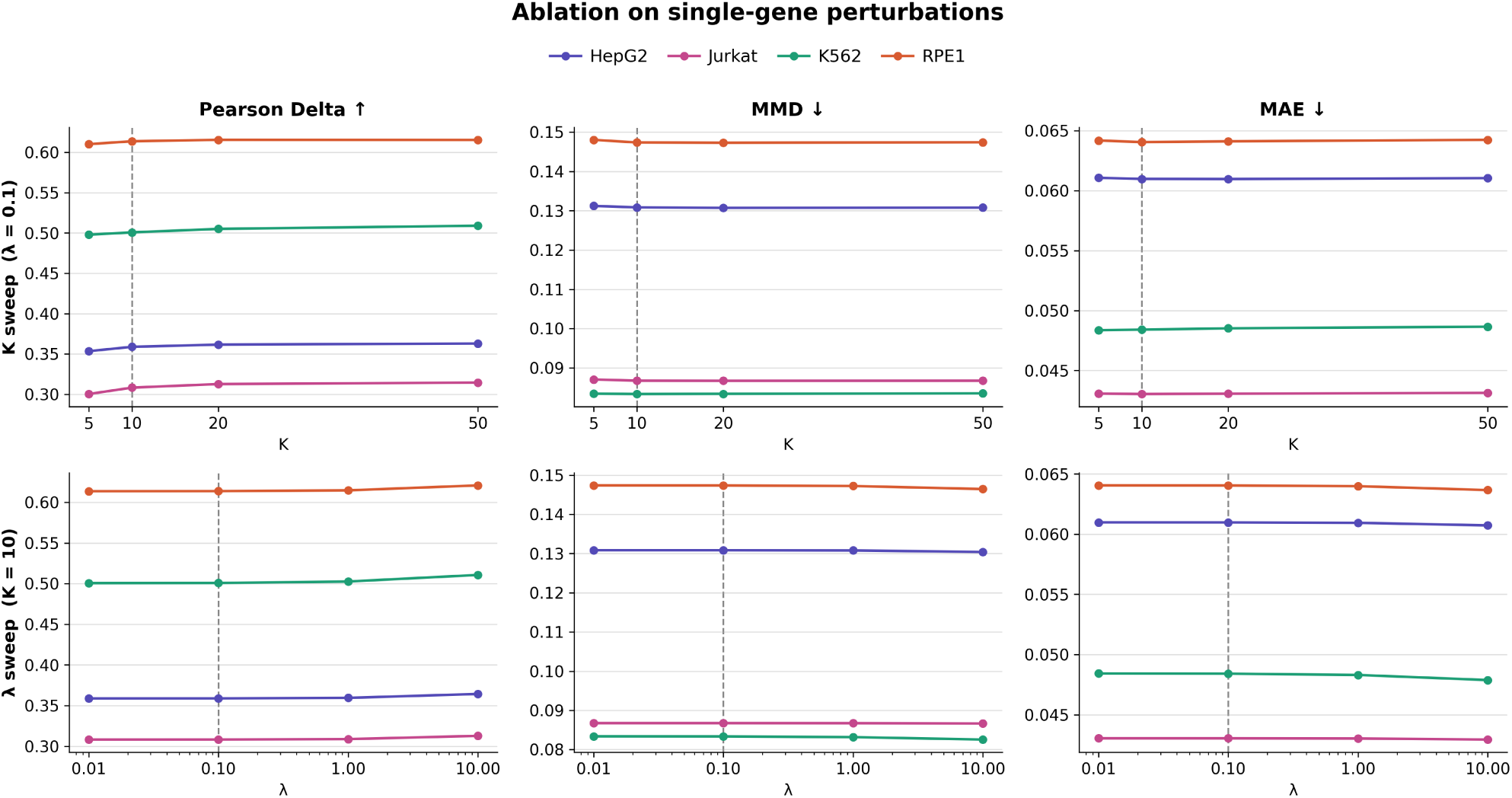
Hyperparameter robustness on single-gene perturbation datasets. Performance of SLIM as a function of the number of PCA components *K* (left column, *λ* = 0.1 fixed) and ridge regularization strength *λ* (right column, *K* = 10 fixed), evaluated on Pearson Delta (higher is better), MMD (lower is better), and MAE (lower is better) across four cell lines. Dashed vertical lines indicate the default setting (*K* = 10, *λ* = 0.1).

**Figure 5.**
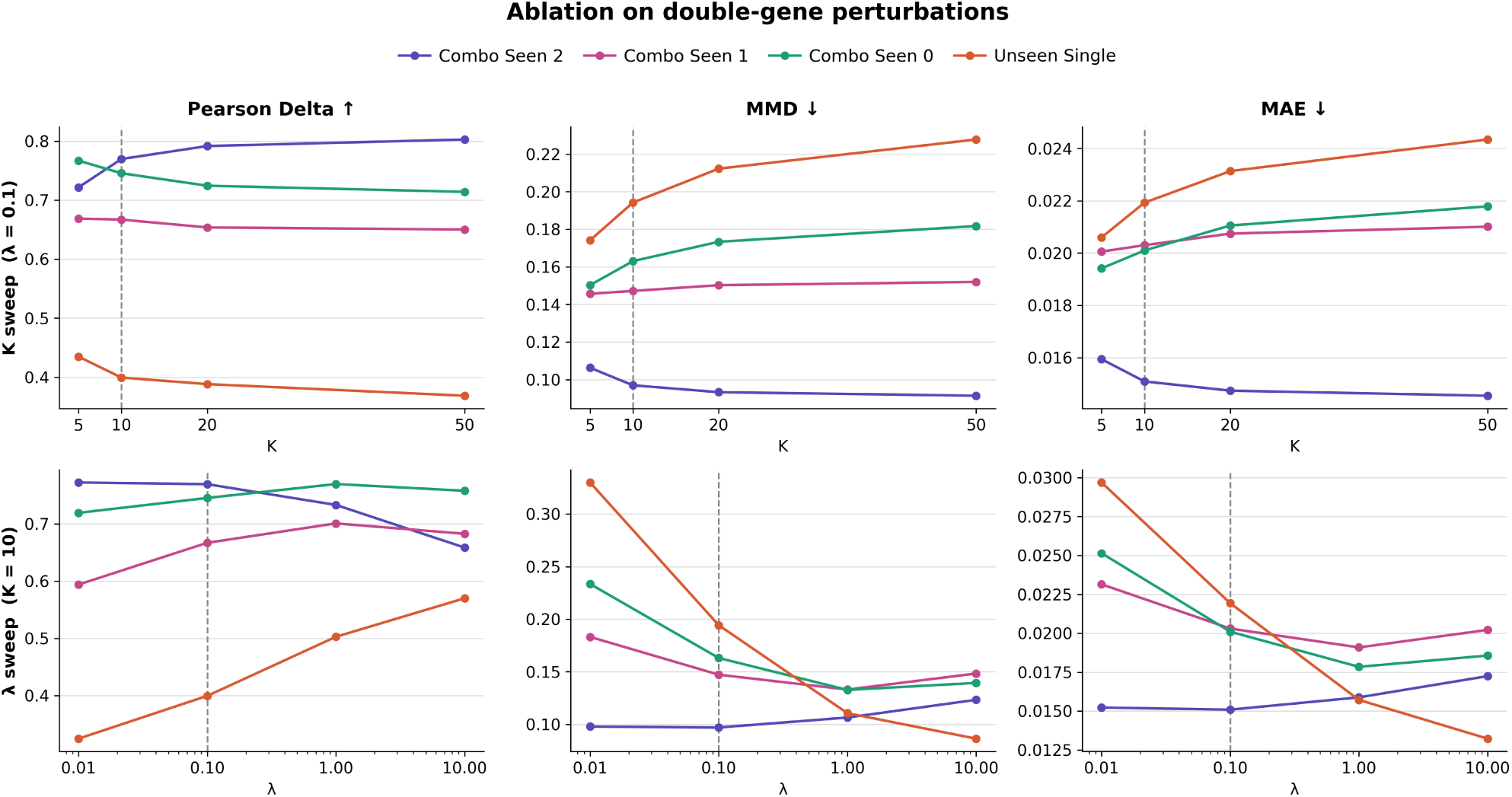
Hyperparameter robustness on the Norman double-gene perturbation dataset. Performance of SLIM as a function of the number of PCA components *K* (left column, *λ* = 0.1 fixed) and ridge regularization strength *λ* (right column, *K* = 10 fixed), evaluated on Pearson Delta (higher is better), MMD (lower is better), and MAE (lower is better) across four splits. Dashed vertical lines indicate the default setting (*K* = 10, *λ* = 0.1).

The Norman dataset showed a more nuanced sensitivity pattern (Figure 5). For unseen single-gene perturbations, increasing *K* progressively degraded all three metrics, whereas stronger regularisation had better scores. Performance on combinatorial perturbations was comparatively stable across both hyperparameters. Training separate models for single- and double-gene perturbations did not resolve this difference (Tables S4 and S5). Instead, the pattern was consistent with overly aggressive extrapolation to unseen genes at the default *λ* = 0.1. With a strong regularisation term *λ* = 10, Pearson Delta on the unseen-single split exceeded the mean baseline. This indicates that the STRING embeddings contained perturbation-specific signal but that its effective use depended on regularisation strength.

To isolate the contribution of the perturbation representation, we replaced the STRING embeddings with two alternatives while leaving the remainder of SLIM unchanged: DepMap embeddings derived from MORPH [20] and GenePT embeddings [8] (Figures S4 and S5). STRING generally matched or outperformed both alternatives, with the clearest advantage in the single-gene benchmarks. In the Norman dataset, GenePT performed similarly to STRING on the combinatorial splits and better on the unseen-single split. These results support the value of prior-informed perturbation embeddings while also showing that no single representation was optimal in every generalisation setting. The per-perturbation distributions are shown in Figures S12 and S13.

### SLIM was orders of magnitude faster than deep learning alternatives

A key practical advantage of SLIM is its computational efficiency. Because the model has a closed-form solution via ridge regression, training requires only two matrix operations: a pseudoinverse in the PCA gene space and a pseudoinverse in the STRING embedding space, each of dimension at most *K* × *D* = 10 × 64. On all five benchmarked datasets, training completed in under 10 seconds on a standard CPU, compared to minutes to hours for GEARS, scGPT, CPA, and scLAMBDA, which require GPU-accelerated iterative optimisation over millions of parameters (Table 1).

**Table 1.** Approximate trainable parameter counts. Trainable parameter counts for SLIM and all benchmarked models. SLIM’s count is the weight matrix W *∈* ℝ^10×64^; its fixed bias and PCA basis are not counted. CPA’s count varies across datasets due to its per-perturbation learned embedding table.

| Model | Trainable parameters (approx.) |
| --- | --- |
| SLIM | 640 |
| GEARS | $\sim 2.3\text{M}$ |
| CPA | $\sim 5.7\text{--}6.7\text{M}$ |
| scLAMBDA | $\sim 9.7\text{M}$ |
| scGPT | $\sim 51.9\text{M}$ |

SLIM’s only trainable parameters were the weight matrix **W** ∈ ℝ^10×64^, just 640 values. The bias vector 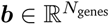 was non-learnable and fixed to the mean expression across training perturbations, and the PCA basis **G** was a fixed data-derived basis rather than an optimised parameter. These 640 trainable parameters were roughly 3,600 times fewer than GEARS (∼2.3M) and roughly 81,000 times fewer than scGPT (∼51.9M).

A practical caveat concerns memory. In our implementation, peak resident memory ranged from 14.7 GB (Norman, 8.9 × 10^4^ cells) to 35.6 GB (Jurkat, 2.6 × 10^5^ cells), scaling approximately linearly with cell number at roughly 150 KB per cell. This cost, however, is almost entirely attributable to the data format rather than to the model: loading the expression matrix and preparing the perturbation splits accounted for 10.5–26.1 GB, and materializing the predicted single-cell populations for evaluation accounted for most of the remainder. The estimator itself is negligible by comparison. Fitting consumes only per-perturbation pseudobulk means (5,000 × 1,428 ≈ 57 MB for Jurkat) and added less than 0.2 GB above the loaded dataset, with fewer than 10^5^ learned parameters occupying under 1 MB. Because the fit depends on the data only through per-perturbation means, memory is in principle reducible to O(*n*_genes_×*n*_perts_) by accumulating those means in a streaming pass, so application to substantially larger screens would require chunked loading rather than any change to the model. Prediction is a single matrix–vector product requiring no GPU and completing in under 0.1 s; the remaining memory cost at prediction time is the elementwise rescaling of pre-sampled control cells, which is a property of the cell-level output format shared by all methods evaluated here.

## Discussion

We present SLIM, an extension of the bilinear framework of Ahlmann-Eltze et al. [9] that combines two components: STRING-based representations of perturbed genes and a retrieval-based procedure that converts predicted mean profiles into cell-level outputs. This design deliberately separates prediction of the perturbation mean from reconstruction of a cell population.

SLIM had a median rank of one across the evaluated dataset–metric combinations and within individual held-out perturbations, and it fitted each benchmark dataset in under 10 seconds on a standard CPU. Its performance was competitive with the strongest evaluated deep learning model despite having only 640 trainable parameters. STRING embeddings offer a compact summary of functional relationships supported by genomic context, high-throughput experiments, conserved co-expression, text mining, and curated databases, rather than relying exclusively on co-expression in particular cellular contexts. The embedding ablation suggests that this information is useful. However, the advantage was not uniform, as GenePT performed better on the Norman unseen-single split.

The largest numerical margin was observed for MMD. This metric asks whether a model reproduces the distribution of the cell population rather than only its mean. Capturing that distribution appears to be difficult: the three deep learning models that generate cell populations natively scored worse on MMD than the linear baseline expanded by independent per-gene sampling, despite comparable mean-response accuracy. We speculate that these models do not explicitly account for population structure, and that learning it from a few hundred training perturbations is a harder estimation problem than the mean-response task they are optimised for. The real training cells, however, already carry this structure, and retrieval with rescaling uses it directly rather than attempting to reconstruct it.

At the dataset scales considered here, the results suggest that the choice of perturbation representation can be at least as consequential as model capacity. This interpretation is consistent with DenAdel et al. [21], who reported diminishing gains from increasing pretraining-set size, and Wang et al. [22], who observed that greater model capacity did not consistently improve Pearson correlation. These findings do not establish biological priors as the sole bottleneck. Cellular-context diversity [23], training objectives [24], preprocessing, and evaluation design may all limit current models.

SLIM has several limitations. First, the model requires perturbation training data for each cell type, and we have not tested whether it can generalize to new cell lines where only control cells are available. We speculate this would be challenging, as the PCA basis and baseline are both fitted to a specific cellular context. This would be expected from the findings in [10], showing that simple linear models perform well when predicting unseen KDs in the same cellular context under low data regimes, but do not excel when predicting across cell types nor when enough fine-tuning data is available. Second, the bilinear structure with a low-rank PCA basis may struggle on more complex datasets where perturbation responses span a higher-dimensional space than ten principal components can capture. This is consistent with the performance drop on the unseen-single split of the Norman dataset, where the correction term extrapolates beyond its training support. Third, SLIM does not generate cell populations from scratch. The retrieval-augmented rescaling strategy borrows heterogeneity from training cells rather than predicting it, meaning perturbation-specific changes in cell-to-cell variability or rare subpopulation responses are out of reach.

Several directions could improve on SLIM’s framework while preserving its simplicity. The PCA representation of readout genes, which is fitted per dataset, could be replaced by dense gene embeddings from a model pre-trained across multiple perturbation datasets, potentially enabling transfer to new cell types without requiring perturbation training data. On the generation side, the retrieval-augmented rescaling strategy could be replaced by a lightweight diffusion model [25] trained to capture the full distributional structure of perturbed cell populations, moving beyond mean-level prediction to faithfully reconstruct perturbation-specific heterogeneity. While we used STRING network embeddings throughout this study, our ablation showed that they generally outperform or match alternative representations. A natural extension would be to combine multiple sources of prior knowledge into a joint perturbation representation.

In summary, SLIM shows that a lightweight linear model informed by protein-network embeddings can provide competitive within-context perturbation predictions with minimal computational cost. The results motivate further work on biological representations and on fair evaluation of the procedures used to reconstruct cell populations. They also caution against attributing performance gains to model capacity alone when training perturbations are limited.

## Methods

### SLIM

#### Model structure

SLIM predicts the mean transcriptional response to a genetic perturbation via a bilinear model, as introduced in [9]. Training data consist of pseudobulked single-cell transcriptomic profiles: gene expression is averaged across all cells belonging to the same perturbation condition to yield one profile per condition. These profiles are stored as columns of 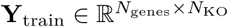, where *N*_genes_ is the number of readout genes and *N*_KO_ is the number of training knockouts.

A perturbation-independent baseline 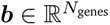 is defined as the mean expression profile across all training perturbations:

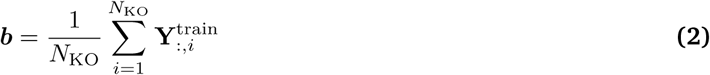

For double-gene perturbation datasets, two separate baselines are maintained: ***b***_single_, the mean expression profile across single-gene training perturbations, and ***b***_double_, the mean across double-gene training perturbations. The appropriate baseline is selected based on the order of the prediction target. This distinction accounts for the dosage effect of simultaneous perturbations [18, 17]: the global expression level under a two-gene perturbation systematically differs from that of a single-gene perturbation, and using a matched baseline absorbs this global offset before fitting **W**.

Readout genes are represented as *K*-dimensional vectors stored in the rows of 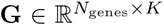. The matrix **G** is obtained by applying PCA to the baseline-centered matrix **Y**_train_ − ***b*** along the perturbation axis, projecting from *N*_KO_ dimensions down to *K*. This representation captures the co-expression structure of readout genes across perturbations in a given cellular context.

Perturbed genes are represented as 64-dimensional vectors stored in the rows of 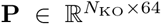. Each perturbed gene is described by a protein-level network embedding derived from the STRING database (v12.0) [14] using the node2vec algorithm [19], as described in [16]. We converted the STRING IDs in the original embedding file into the IDs used in the benchmark datasets via the STRING ID mapping API (https://string-db.org/help/api). After ID mapping, all the perturbations in the used benchmarks were covered. These embeddings encode functional similarities between proteins based on experimental evidence, text mining, and curated pathway knowledge and serve as biology-informed priors for unseen perturbations. For single-gene perturbations, we directly used their STRING embeddings, while for double-gene perturbations, we used their averaged embeddings.

The bilinear map **W** ∈ ℝ*^K^*^×^*^D^* connects the readout gene space to the perturbed gene space. The full model is:

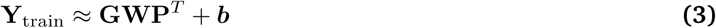

Model parameters **W** are obtained by minimizing the squared reconstruction error:

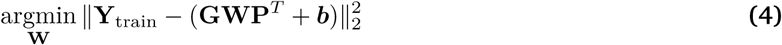

This objective function has a closed-form solution via:

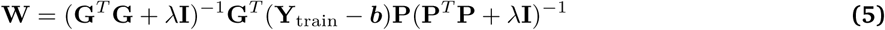

where *λ* is the ridge regularization coefficient applied symmetrically to both the gene-space and perturbation-space factors, and **I** denotes identity matrices of the appropriate size.

#### Hyperparameters

SLIM has two hyperparameters: the number of principal components *K* and the ridge regularization strength *λ*. The embedding dimensionality *D* is fixed by the STRING embeddings and is not tuned. Based on systematic experiments across all benchmarked datasets, *K* = 10 and *λ* = 0.1 yield consistently robust performance across a range of datasets. These values are fixed for all reported results without dataset-specific tuning.

#### Inference

Given unseen perturbation genes *g̃*_1_*, …, g̃_M_*, SLIM retrieves their STRING embeddings to form **P̃**∈ ℝ*^M^*^×^*^D^* and predicts the mean expression profile for each perturbation:

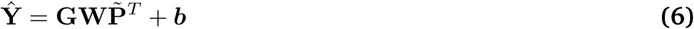

To construct a population of single-cell outputs from a predicted mean, SLIM employs retrieval-augmented rescaling. For each test perturbation, *n* cells are sampled uniformly from the full training pool, where *n* matches the number of cells observed for that perturbation in the test set. Sampling is performed with replacement when *n* exceeds the pool size. In the combinatorial dataset, cells for single- and double-gene predictions are sampled from the corresponding single- and double-perturbation training pools, respectively. Each sampled cell 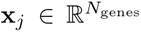 is then multiplicatively rescaled so that the population mean matches the predicted mean 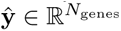:

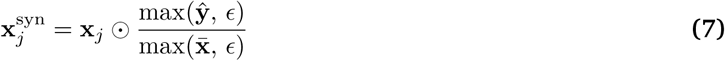

where 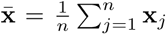 is the per-gene mean of the sampled cells, ⊙ denotes element-wise multiplication, and *ɛ* = 10^−6^ guards against division by zero. This strategy preserves the single-cell heterogeneity structure from the training data while shifting the population mean to the model’s prediction.

### Benchmark datasets

We evaluated SLIM on five perturbation datasets spanning diverse cell types and experimental designs. For single-gene perturbation benchmarks, we used four datasets: K562 chronic myelogenous leukemia cells and RPE1 retinal pigment epithelium cells from the Replogle et al. genome-wide essential gene screen [2], as well as HepG2 hepatocellular carcinoma cells and Jurkat T lymphocyte cells [17]. For the double-gene perturbation benchmark, we used the Norman dataset [18], which contains combinatorial gene overexpression perturbations in K562 cells. All datasets were loaded and partitioned using the GEARS framework [5] via the simulation split, in which all test perturbations are withheld entirely from training.

### Models for comparison

We benchmarked SLIM against four deep learning models: scLAMBDA [7] (v1.0.1), GEARS [5] (cell-gears v0.1.2), scGPT [6] (scgpt v0.2.4), and CPA [4] (cpa-tools v0.8.8). Each model was run in a dedicated conda environment. scLAMBDA was run from a source checkout of the authors’ official implementation, as it is not distributed on PyPI. We also included the linear model of Ahlmann-Eltze et al. [9], which suggested the **GWP***^T^* + ***b*** formulation but used the same PCA-derived representations for **G** and **P**^T^.

Methods that natively generate single-cell populations (scLAMBDA, scGPT, and CPA) were used as-is. For methods that produce only mean expression profiles (linear model and GEARS), we generated synthetic cell populations by drawing each gene independently from a Gaussian parameterized by the predicted mean and the per-gene standard deviation computed from the training set. To study the benefit of our random sampling and rescaling strategy, we added SLIM-Gaussian, which uses the same predicted mean profile as SLIM, but expands to the cell population by the per-gene standard deviation as linear model and GEARS. For all methods, the number of synthetic cells per perturbation was set equal to the number of ground-truth test cells for that perturbation.

### Evaluation metrics

We evaluated all methods using three metrics computed against the ground-truth test cells for each perturbation. For all three metrics, the per-perturbation metrics are averaged across all test perturbations to obtain a single score per method and dataset. Meanwhile, we reported the per-perturbation score distributions in the supplementary material.

#### Pearson Delta

For each perturbation *p*, let 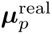 and 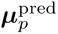 denote the mean expression profiles of the real and predicted cells, and let ***µ***_ctrl_ denote the control means. The differential expression vectors are 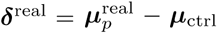 and 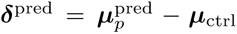. Subtracting the control removes the shared baseline transcriptome, which would otherwise dominate the correlation. Pearson Delta is the Pearson correlation between ***δ***^real^ and ***δ***^pred^ across genes:

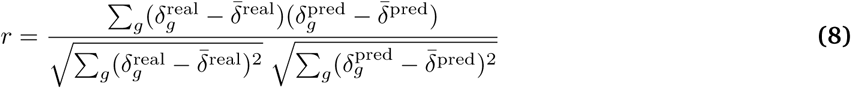

Higher Pearson Delta indicates that the model better reproduces which genes respond and how strongly they respond relative to one another. Note that *r* is invariant to positive affine transformations of ***δ***^pred^: a prediction that recovers the correct response pattern but systematically underestimates its overall amplitude (***δ***^pred^ = *a* ***δ***^real^, *a >* 0) attains *r* = 1 regardless of *a*. Pearson Delta therefore measures the shape of the perturbation response, and we report MAE and MMD alongside it to assess absolute magnitude and distributional agreement.

#### Maximum Mean Discrepancy

Maximum mean discrepancy (MMD) [26] measures the discrepancy between the full predicted and real single-cell distributions. Given real cells 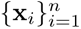 and predicted cells 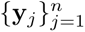 of equal size (at most 500 cells), MMD is estimated with a mixture of *T* = 5 Gaussian kernels:

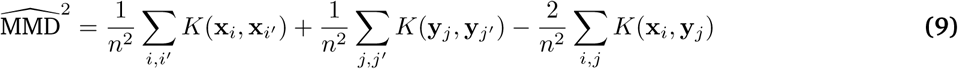

where 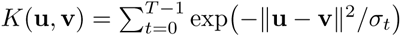 and the bandwidths *σ_t_* = *σ*_0_·2*^t^* are set by the median heuristic, with *σ*_0_ equal to the median pairwise squared distance divided by 2^⌊^*^T/^*^2⌋^. Lower MMD indicates that the predicted cell population is more similar to the real population in distribution.

#### Mean Absolute Error

The mean absolute error (MAE) is the average absolute difference between the predicted and observed mean expression profiles:

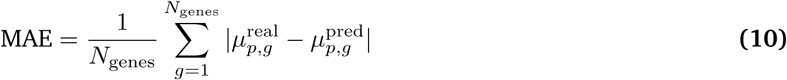

Lower MAE indicates closer agreement between the predicted and observed mean expression profiles.

## Data and code availability

Source code is available at https://github.com/RasmussenLab/SLIM. The detailed scores and plots are available at https://doi.org/10.5281/zenodo.21837272. The benchmark datasets are third-party datasets described and cited in the Benchmark datasets section.

## Acknowledgements

We would like to thank the other members of the Rasmussen lab, in particular Foteini Aktypi, for our fruitful discussions regarding the project. The authors used CodeX and Claude to improve the readability and language of the manuscript. No content, results, or interpretations were generated by AI. All authors reviewed and edited the output and take full responsibility for the content of the publication.

## Supplementary materials

**Table S1.** Number of perturbations in each single-perturbation dataset split. While SLIM covers all the perturbations in these datasets, values in parentheses indicate perturbations covered by all benchmarked models.

| Dataset | Train | Validation | Test |
| --- | --- | --- | --- |
| K562 | 733 | 82 | 272 (270) |
| RPE1 | 1035 | 115 | 384 (375) |
| Jurkat | 1428 | 161 | 535 (515) |
| HepG2 | 1217 | 137 | 455 (435) |

**Table S2.** Number of perturbations in the Norman double-perturbation dataset. Single and double refer to the number of genes perturbed simultaneously. While SLIM covers all the perturbations in these datasets, values in parentheses indicate perturbations that are covered by all benchmarked models.

| Perturbation type | Train | Validation | Test |
| --- | --- | --- | --- |
| Single | 99 | 13 | 26 (25) |
| Double | 39 | 18 | 71 (69) |
| Total | 138 | 31 | 97 (94) |

**Table S3.** Number of test perturbations in the Norman double-perturbation dataset by split category. The dataset defines four splits of increasing difficulty based on how many of the two perturbed genes were seen as single perturbations during training: combo seen 2 (both genes seen individually), combo seen 1 (one seen), combo seen 0 (neither seen), and unseen single (a single-gene perturbation held out entirely). While SLIM covers all the perturbations, values in parentheses indicate perturbations covered by all benchmarked models.

| Test split | Perturbations |
| --- | --- |
| Unseen single | 26 (25) |
| Combo seen 2 | 19 (19) |
| Combo seen 1 | 43 (42) |
| Combo seen 0 | 9 (8) |
| Total | 97 (94) |

**Table S4.** Separate single-perturbation model on the Norman unseen-single split. Performance across *λ* values with *K* = 10 fixed. The model was trained on single perturbations only.

| $\lambda$ | Pearson Delta ↑ | MSE ↓ | MAE ↓ |
| --- | --- | --- | --- |
| 0.001 | 0.26 | 0.0226 | 0.0406 |
| 0.01 | 0.32 | 0.0107 | 0.0298 |
| 0.1 | 0.41 | 0.0056 | 0.0220 |
| 1 | 0.52 | 0.0028 | 0.0157 |
| 10 | 0.58 | 0.0022 | 0.0134 |
| 100 | 0.57 | 0.0024 | 0.0136 |

**Table S5.**
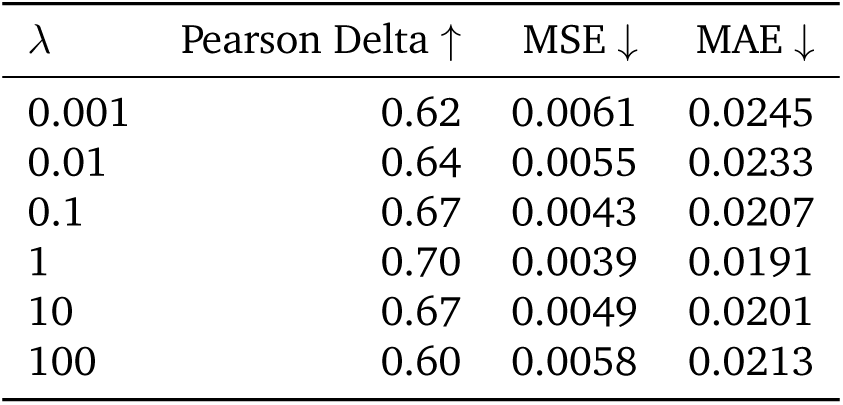
Separate double-perturbation model on the Norman combo splits. Performance across *λ* values with *K* = 10 fixed. The model was trained on double perturbations only.

| $\lambda$ | Pearson Delta ↑ | MSE ↓ | MAE ↓ |
| --- | --- | --- | --- |
| 0.001 | 0.62 | 0.0061 | 0.0245 |
| 0.01 | 0.64 | 0.0055 | 0.0233 |
| 0.1 | 0.67 | 0.0043 | 0.0207 |
| 1 | 0.70 | 0.0039 | 0.0191 |
| 10 | 0.67 | 0.0049 | 0.0201 |
| 100 | 0.60 | 0.0058 | 0.0213 |

**Figure S1.**
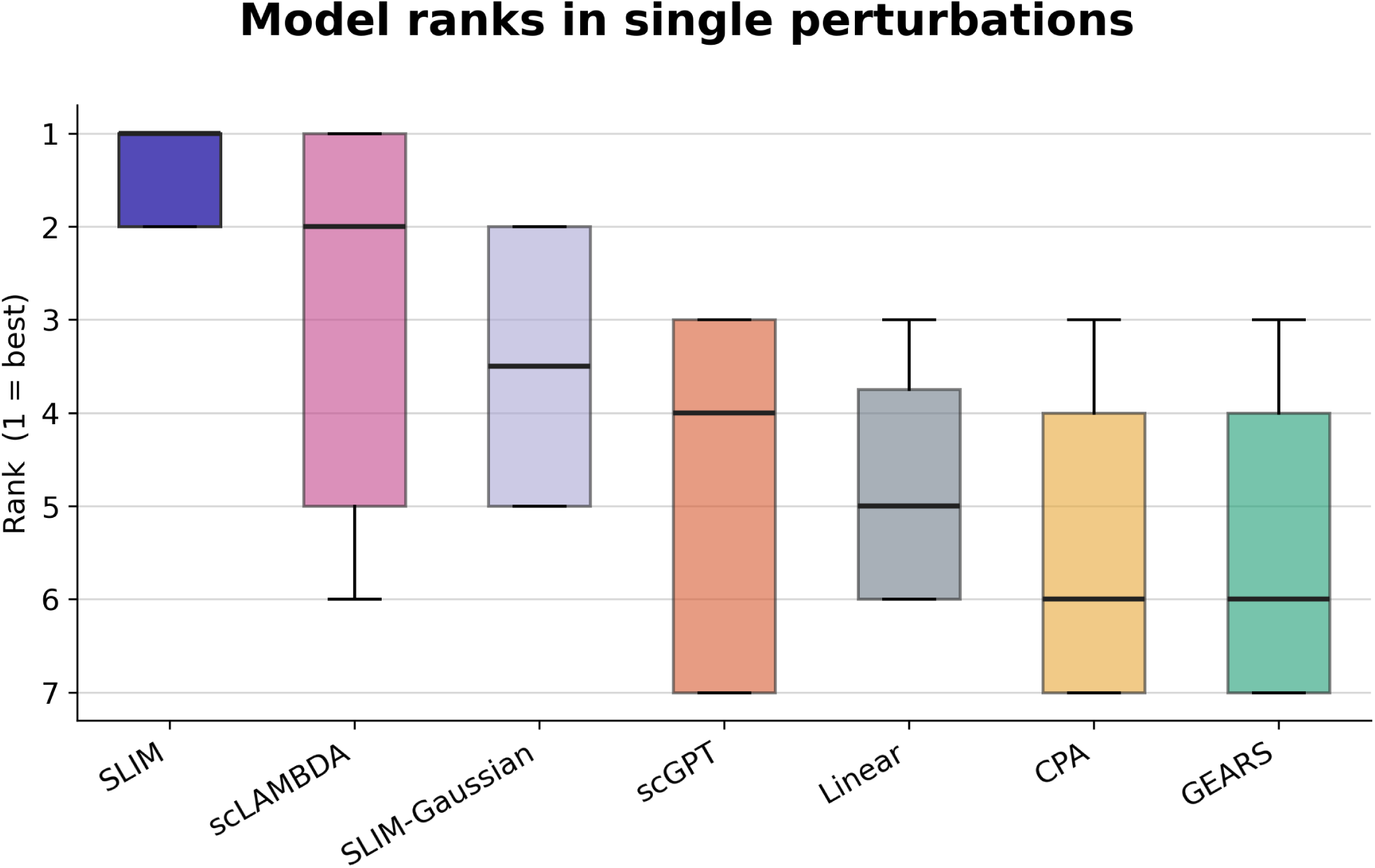
Model rank distribution across single-gene perturbation benchmarks. Rank distributions (1 = best) aggregated across 4 datasets and 3 metrics (12 total rankings). SLIM achieves a median rank of 1, placing first on eight of twelve dataset-metric combinations and second on the remaining four, consistently outperforming all competing methods.

**Figure S2.**
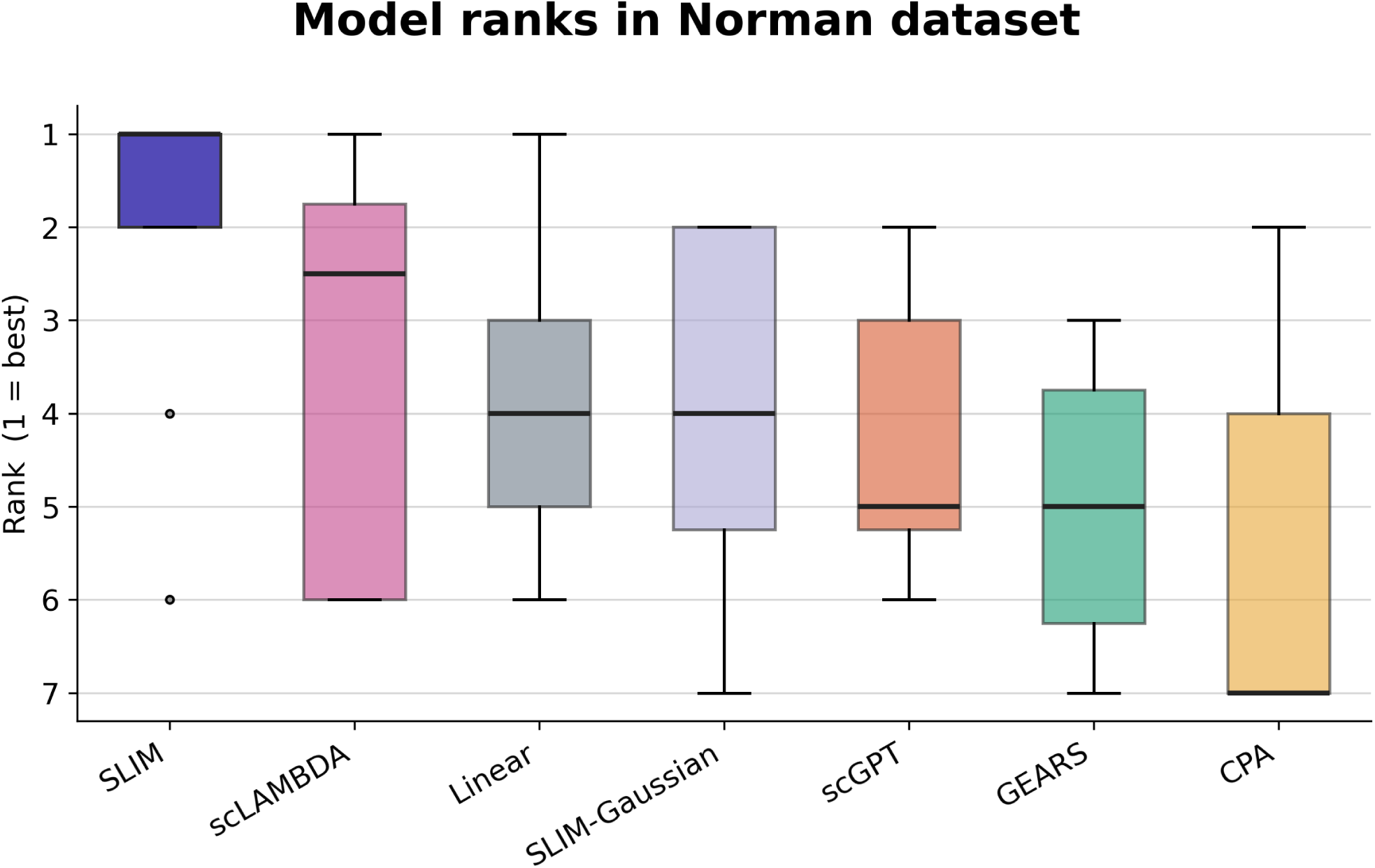
Model rank distribution across double-gene perturbation benchmarks. Rank distributions aggregated across the Norman dataset split groups and 3 metrics. SLIM again ranks first in the large majority of comparisons.

**Figure S3.**
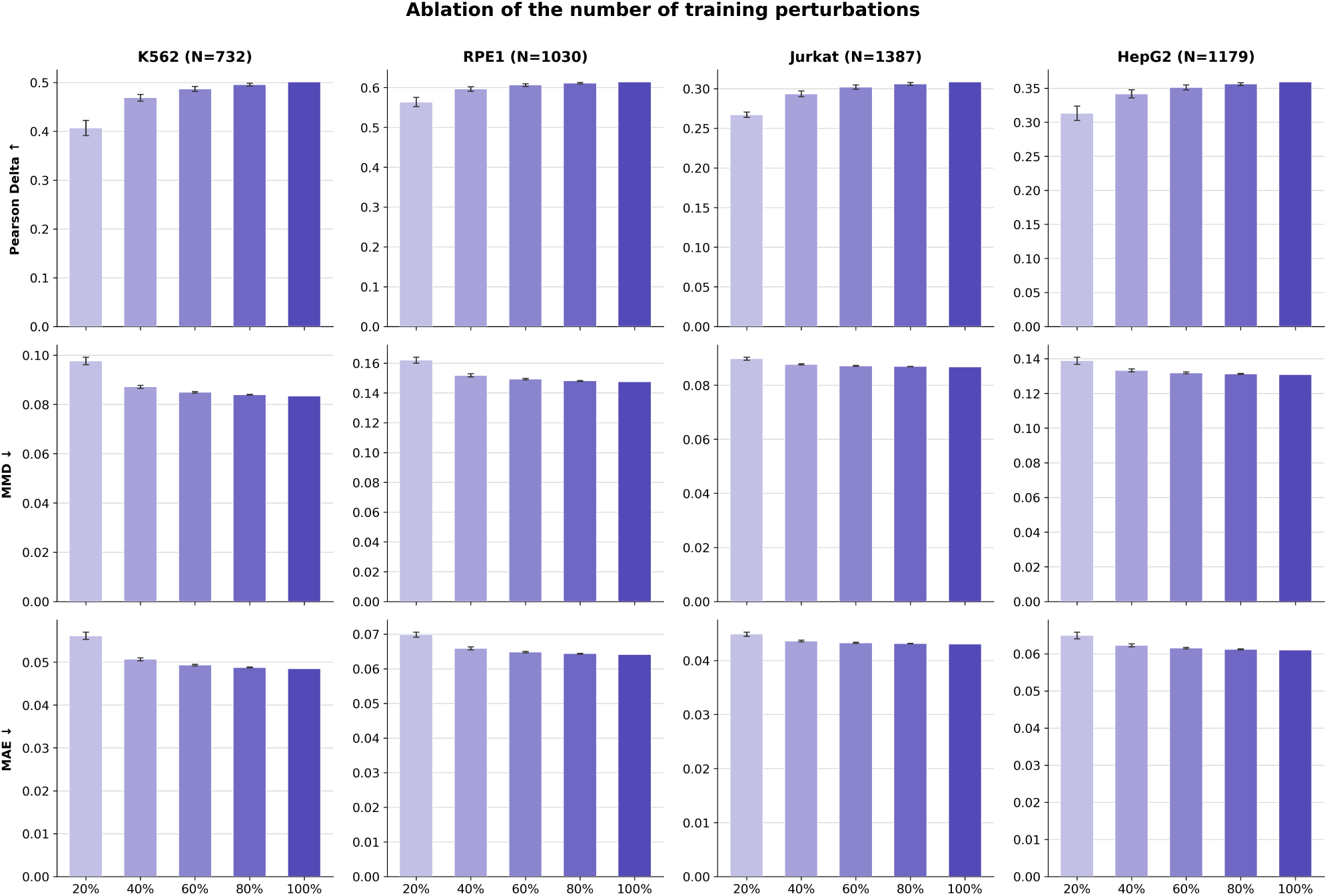
Ablation study on the number of training perturbations in single perturbation datasets. We split the training set of each dataset into five folds and trained SLIM models on any *K* folds and evaluated them on the same test set. The x-axis shows the percentages of training samples, and the total number of perturbations is shown near the dataset names.

**Figure S4.**
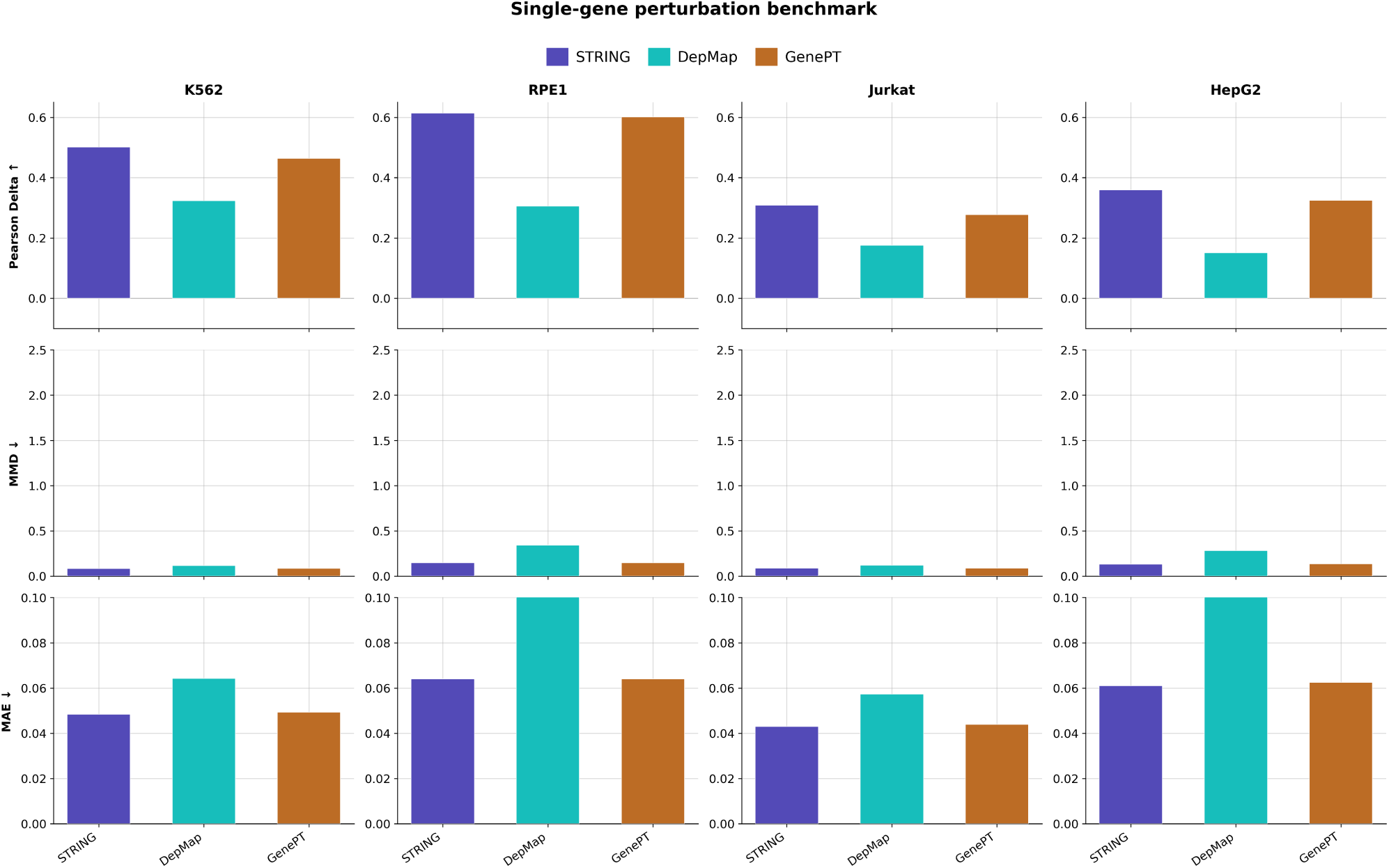
Ablation study on the perturbation embeddings in the single perturbation datasets. We tried the other two gene embeddings in our SLIM framework: DepMap embeddings and GenePT embeddings in single-perturbation datasets. The results show that the STRING embeddings are generally slightly better than GenePT embeddings, and both consistently outperform DepMap embeddings. The DepMap embeddings are sourced from this study [20].

**Figure S5.**
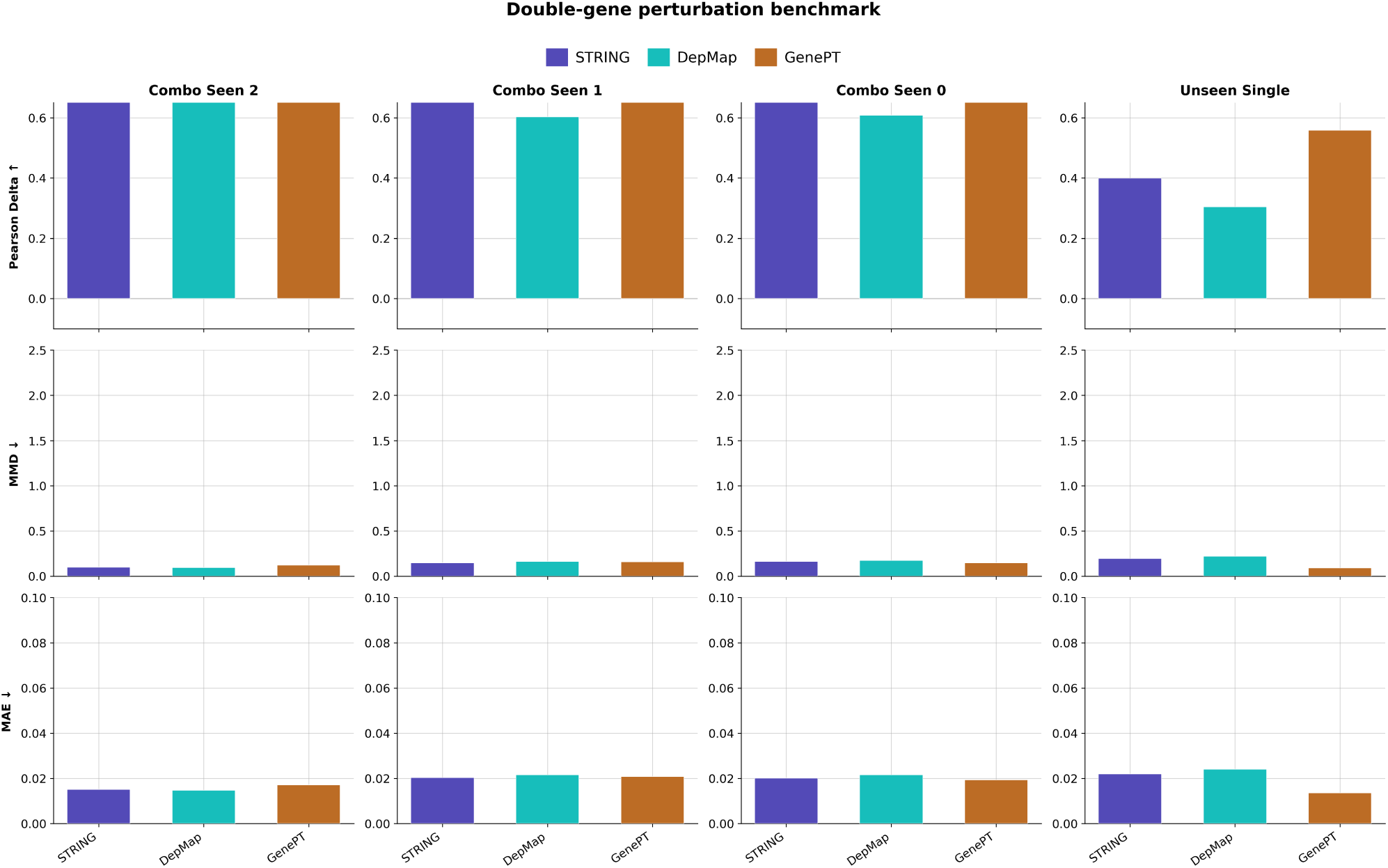
Ablation study on the perturbation embeddings in the double perturbation dataset. We tried the other two gene embeddings in our SLIM framework: DepMap embeddings and GenePT embeddings in the double-perturbation dataset (Norman). The results show that STRING and GenePT have close performance on double perturbations, but GenePT outperformed STRING and DepMap embeddings in single perturbations in this dataset. The DepMap embeddings are sourced from this study [20].

**Figure S6.**
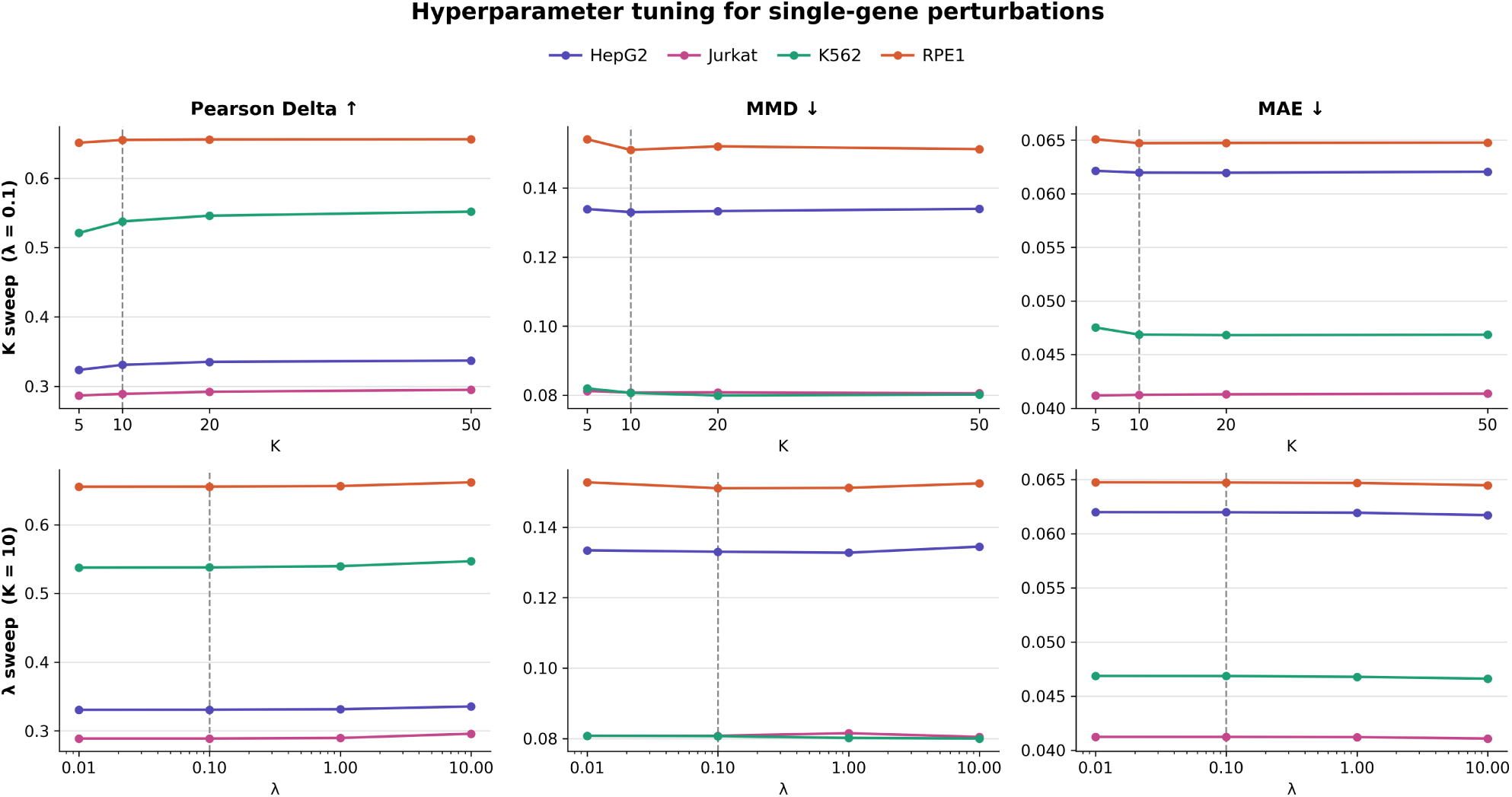
Hyperparameter tuning for single-gene perturbations. Performance of SLIM as a function of the number of PCA components *K* (left column, *λ* = 0.1 fixed) and ridge regularization strength *λ* (right column, *K* = 10 fixed), evaluated on Pearson Delta (higher is better), MMD (lower is better), and MAE (lower is better) across four cell lines. Dashed vertical lines indicate the default setting (*K* = 10, *λ* = 0.1).

**Figure S7.**
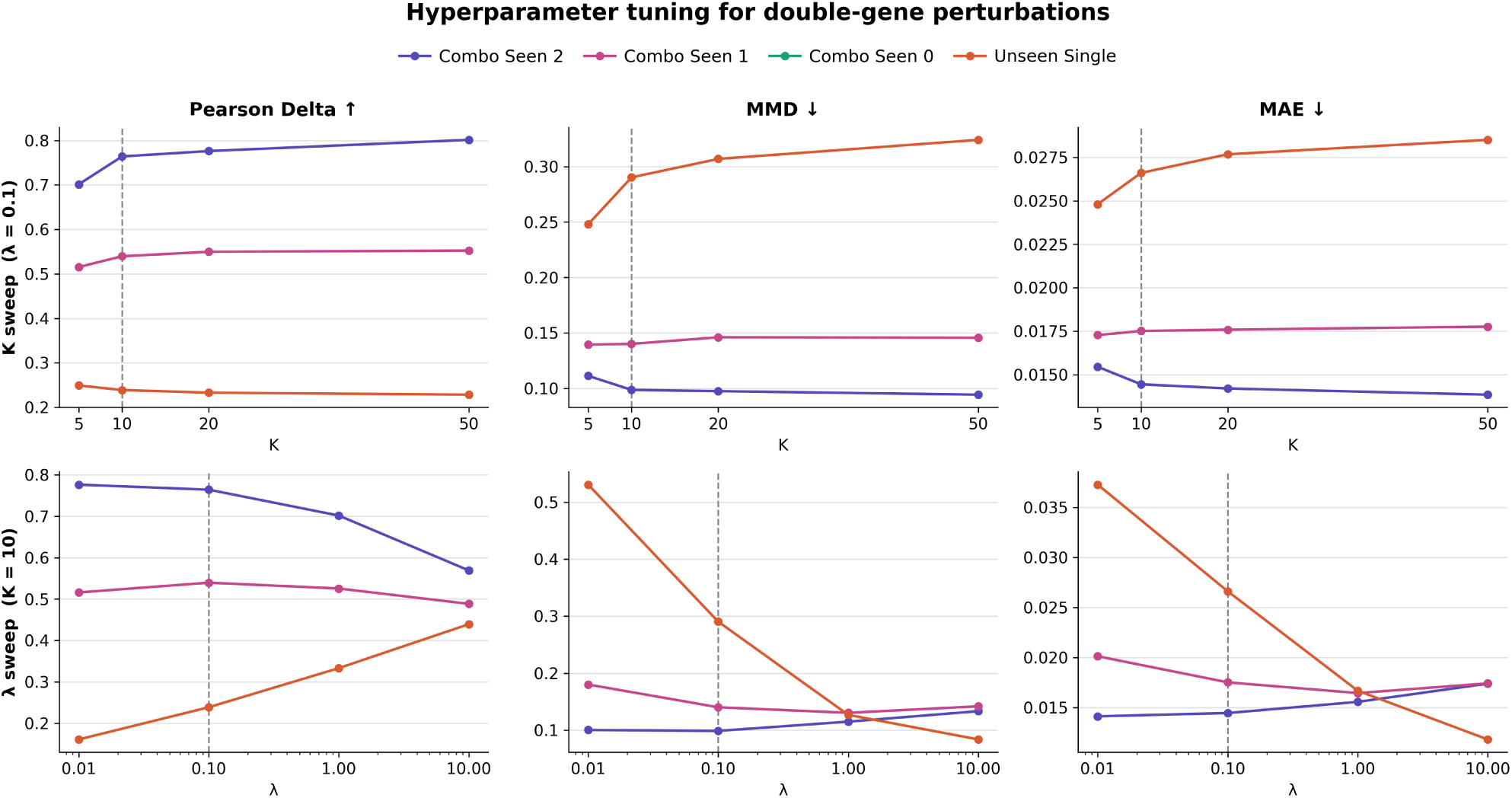
Hyperparameter tuning for double-gene perturbations. Performance of SLIM as a function of the number of PCA components *K* (left column, *λ* = 0.1 fixed) and ridge regularization strength *λ* (right column, *K* = 10 fixed), evaluated on Pearson Delta (higher is better), MMD (lower is better), and MAE (lower is better) across four cell lines. Dashed vertical lines indicate the default setting (*K* = 10, *λ* = 0.1).

**Figure S8.**
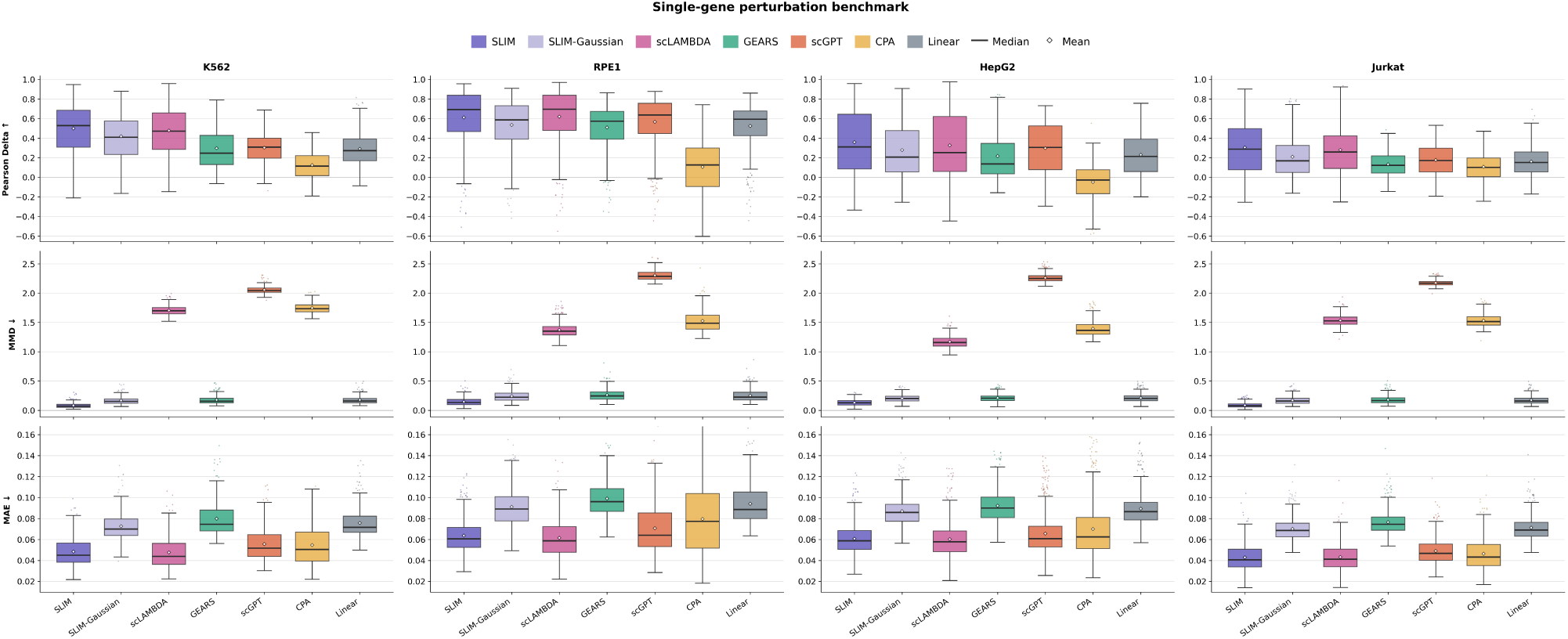
Per-perturbation metric distributions for the single-gene perturbation benchmark. Distribution view of Figure 2: instead of the dataset-level aggregate, each box summarizes the per-perturbation values of Pearson Delta (higher is better), MMD (lower is better), and MAE (lower is better) across the K562, RPE1, HepG2, and Jurkat test sets. Boxes span the interquartile range with the median as a solid line and the mean as a white diamond; whiskers extend to 1.5*×* IQR and outlying perturbations are shown as individual points. Dashed lines indicate the training-perturbation mean and linear baselines, with shaded bands giving their interquartile ranges. The aggregate differences are driven by consistent shifts of the whole distribution rather than by a few perturbations, and the per-perturbation spread is large relative to the gaps between methods on Pearson Delta and MAE, whereas the MMD advantage of SLIM separates the distributions almost completely.

**Figure S9.**
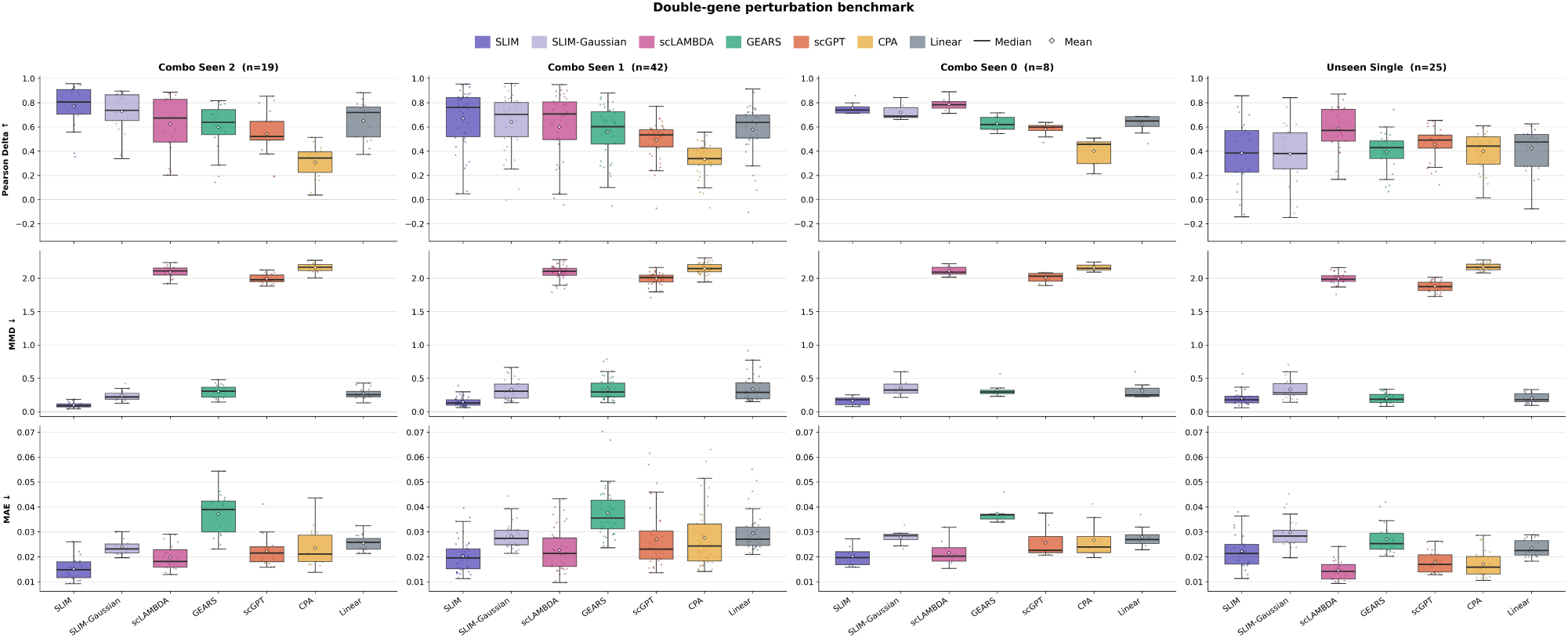
Per-perturbation metric distributions for the double-gene perturbation benchmark. Distribution view of Figure 3 on the Norman dataset, with one panel column per test split (Combo Seen 2, Combo Seen 1, Combo Seen 0, and Unseen Single; the number of test perturbations *n* is given in each panel title). Boxes, medians, means, whiskers, points, and baseline lines follow the conventions of Figure S8. The small number of perturbations in the Combo Seen 0 split, in particular, makes its aggregate values sensitive to individual perturbations.

**Figure S10.**
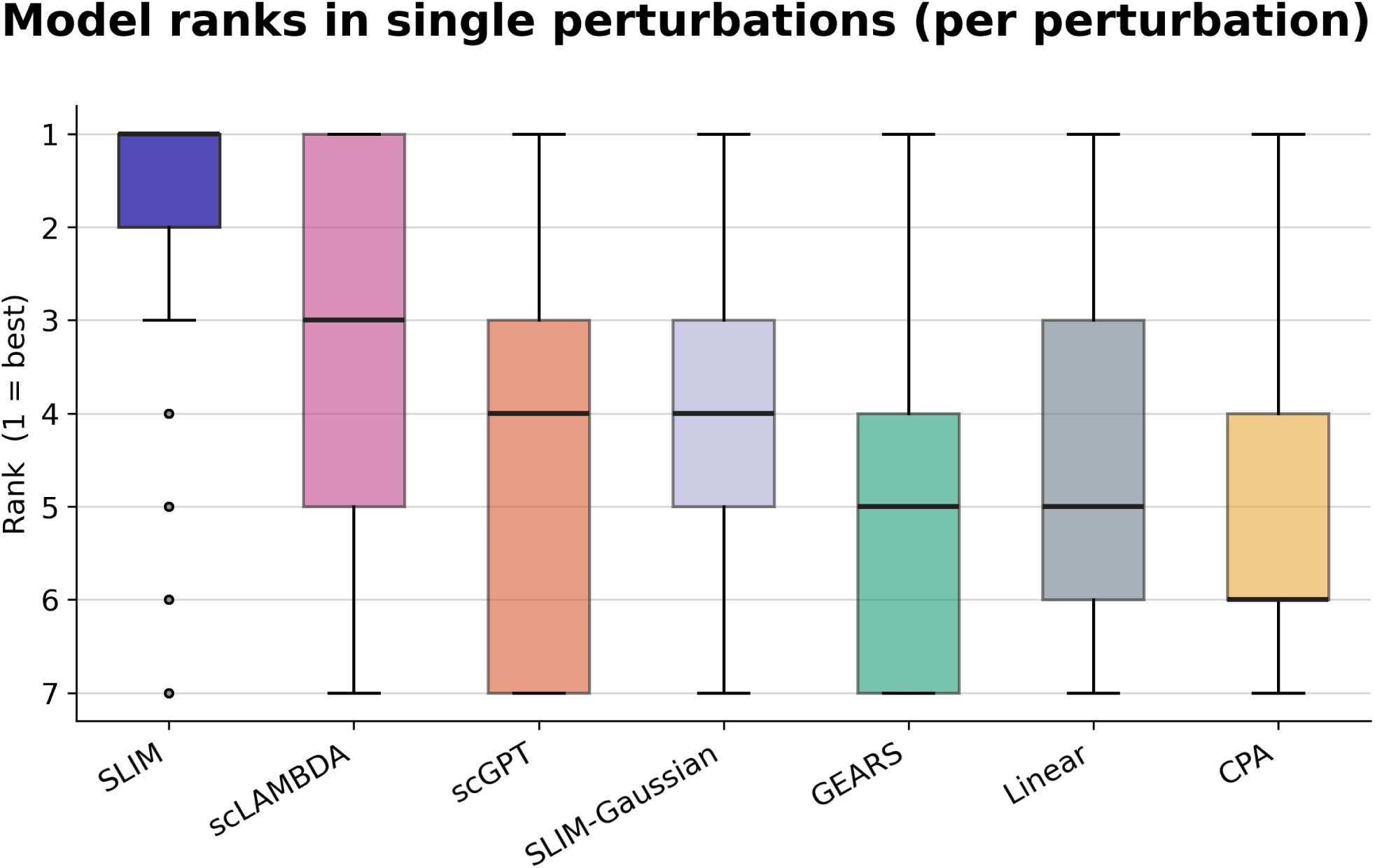
Per-perturbation model rank distribution across single-gene perturbation benchmarks. Complement to Figure S1, which ranks methods on dataset-level aggregates. Here each method is ranked separately within every individual test perturbation (1 = best) against all other methods and the two baselines, and the ranks are pooled across the four single-perturbation datasets and three metrics. SLIM has the best median rank and the tightest rank distribution, showing that its advantage holds perturbation by perturbation rather than only after aggregation.

**Figure S11.**
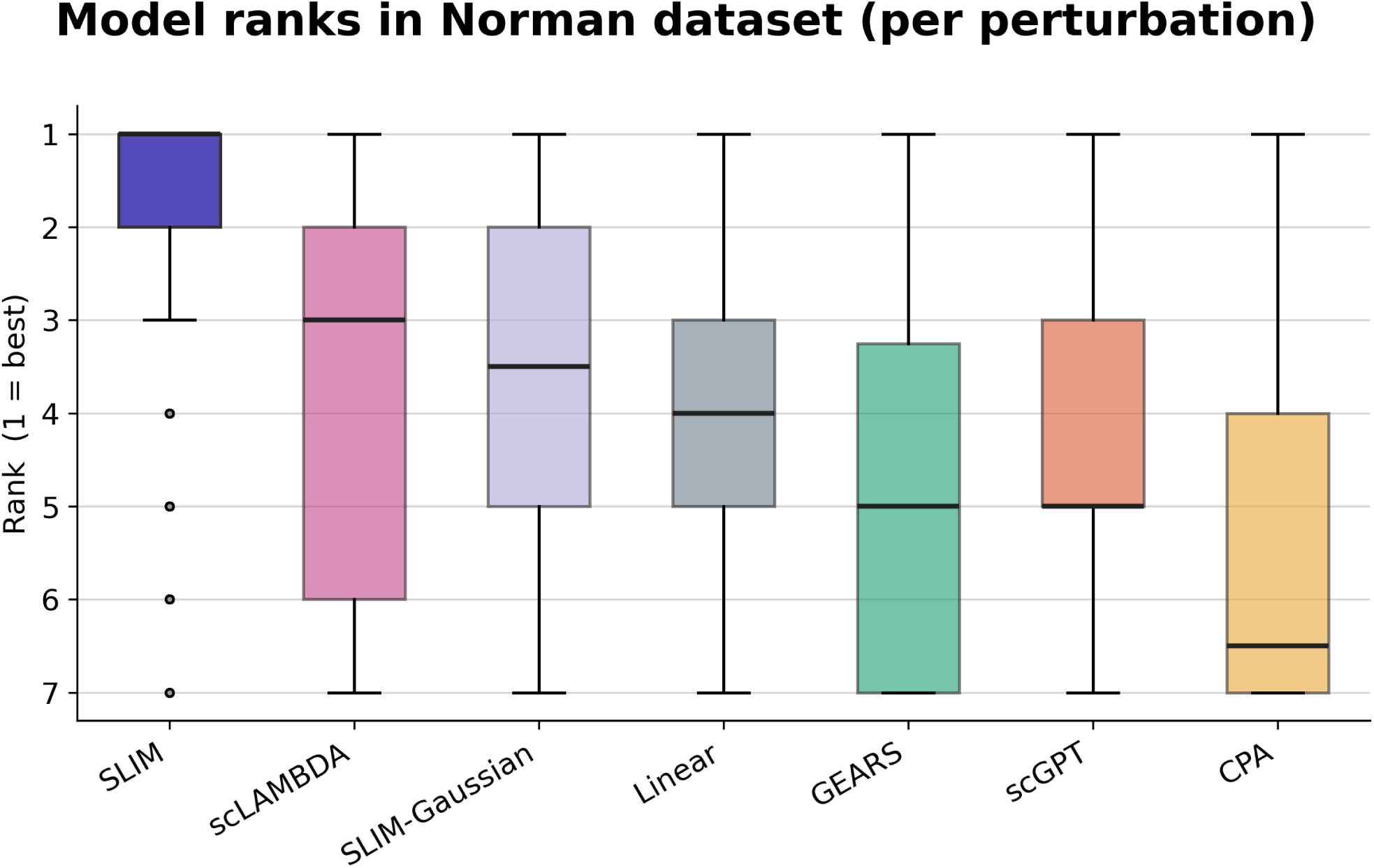
Per-perturbation model rank distribution on the Norman double-gene perturbation dataset. Complement to Figure S2, computed per test perturbation as in Figure S10 and pooled across the Norman split groups and three metrics. SLIM again attains the best median rank, though the spread is wider than in the single-perturbation setting, reflecting the smaller number of test perturbations and the greater difficulty of the combinatorial splits.

**Figure S12.**
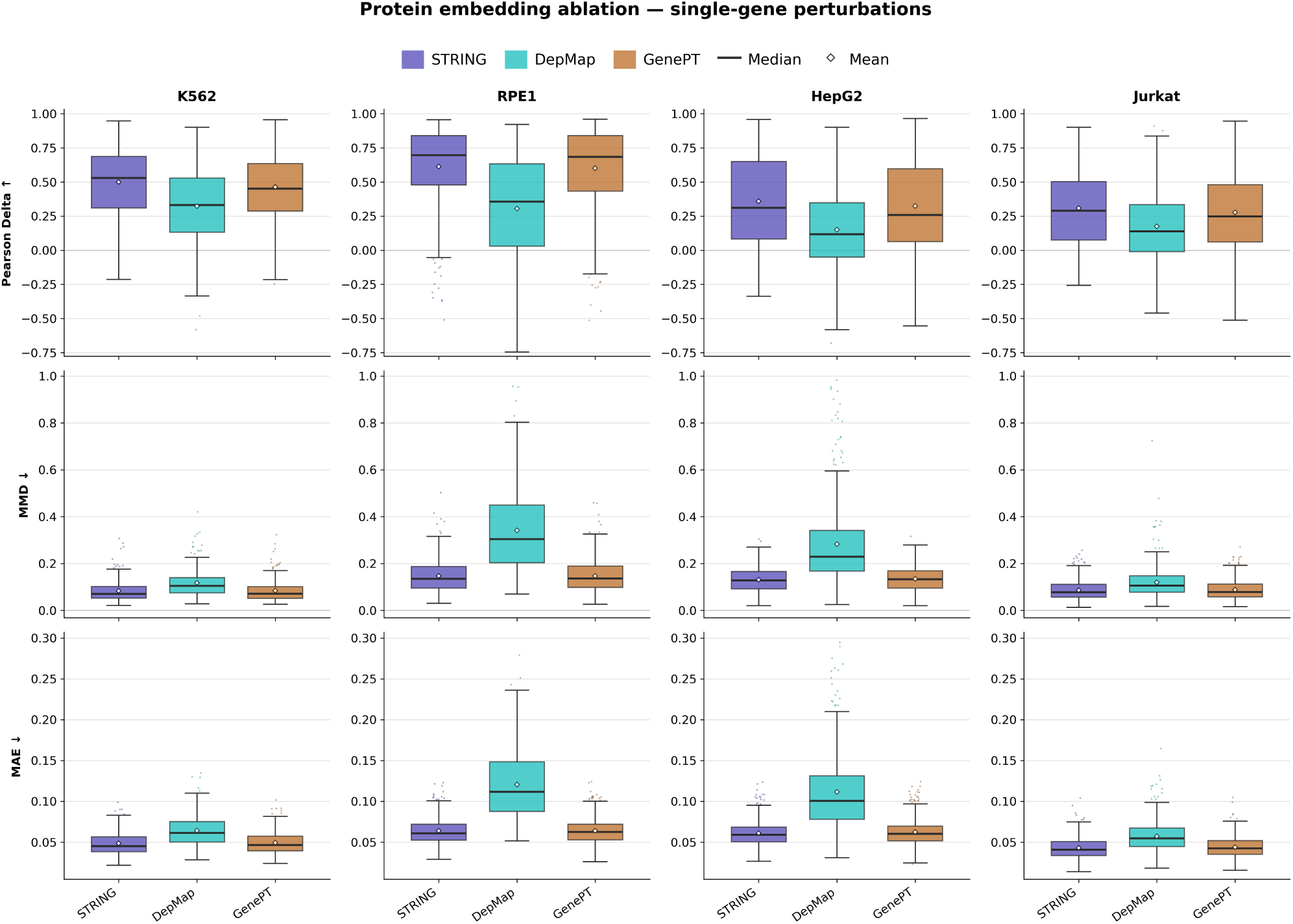
Per-perturbation metric distributions for the perturbation embedding ablation in the single perturbation datasets. Distribution view of Figure S4, comparing STRING, DepMap, and GenePT embeddings within the SLIM framework on the four single-perturbation datasets. Boxes, medians, means, whiskers, and points follow the conventions of Figure S8. STRING and GenePT give closely overlapping distributions on all three metrics, while the DepMap distributions are shifted towards worse values and have noticeably heavier upper tails on MMD and MAE, indicating that the DepMap gap is driven in part by a subset of poorly predicted perturbations.

**Figure S13.**
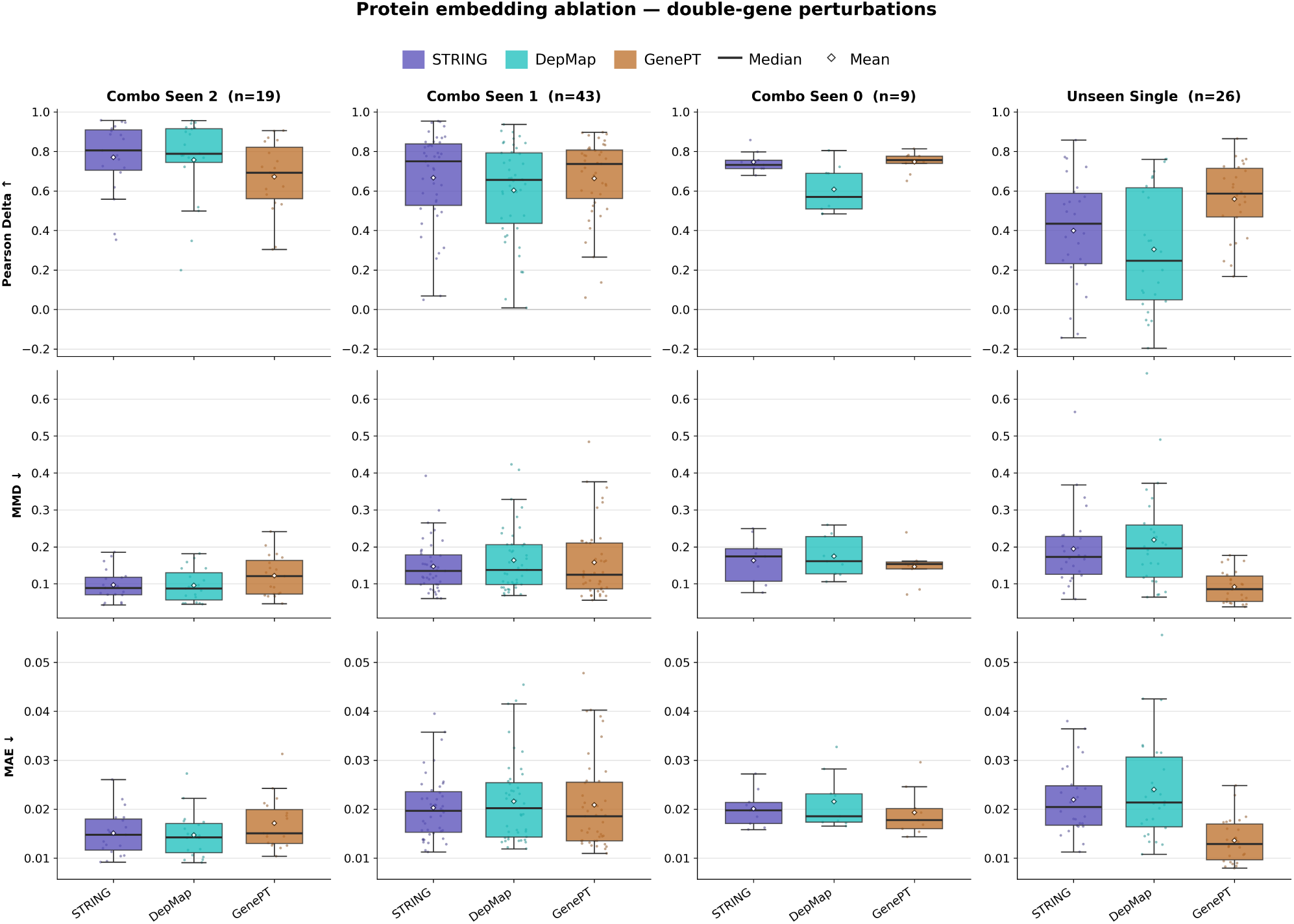
Per-perturbation metric distributions for the perturbation embedding ablation in the double perturbation dataset. Distribution view of Figure S5 on the Norman dataset, split by test category as in Figure S9. The three embeddings produce largely overlapping per-perturbation distributions on the combo splits; the advantage of GenePT reported for this dataset is concentrated in the unseen-single split, where its distribution is shifted towards higher Pearson Delta and lower MMD and MAE. Given the small number of test perturbations per split, these differences should be read as suggestive rather than conclusive.

**Figure S14.**
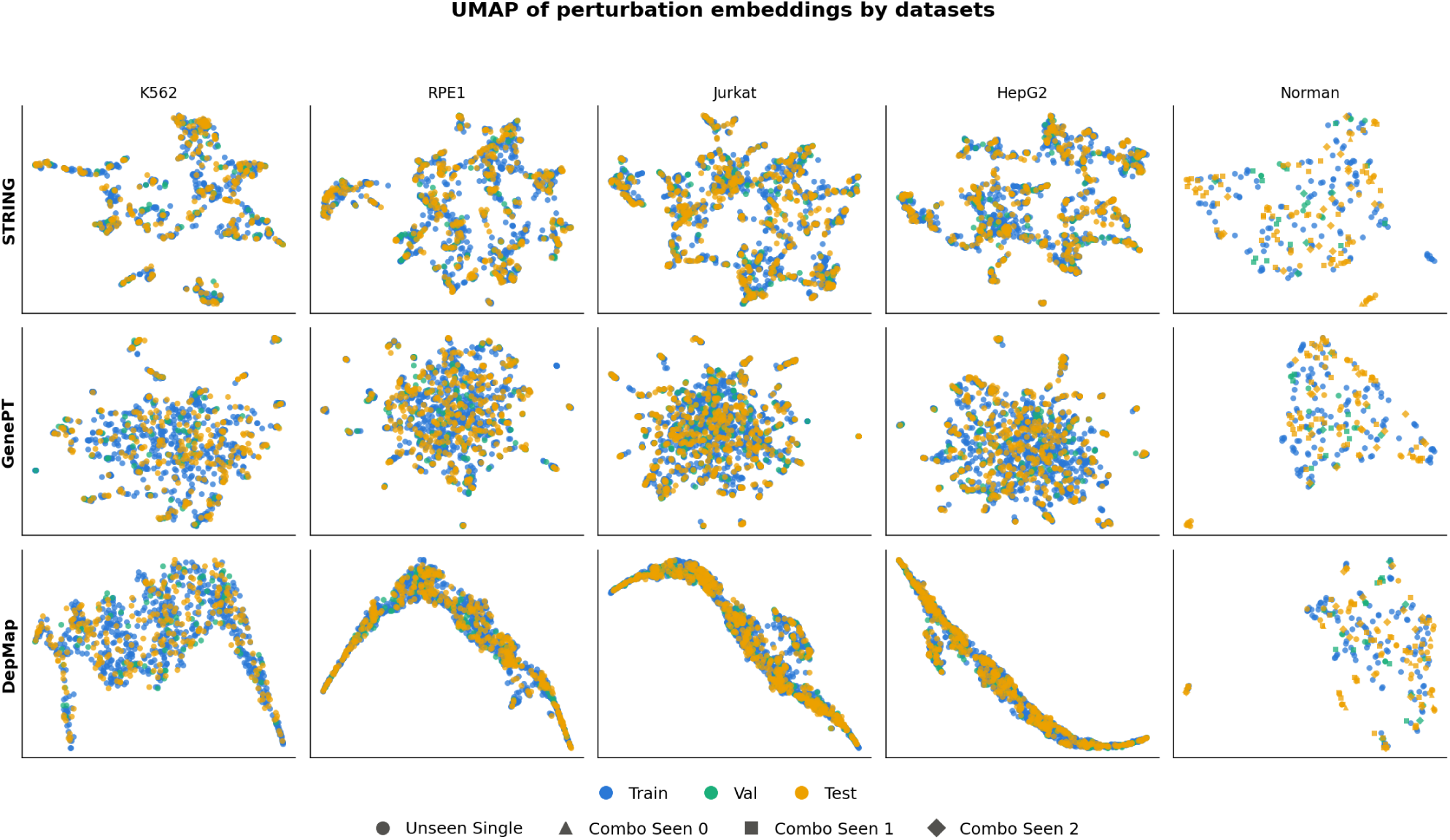
UMAP of perturbation embeddings by datasets. Two-dimensional UMAP projections of perturbation embedding vectors for all five datasets (columns: K562, RPE1, Jurkat, HepG2, Norman) under three embedding methods (rows: STRING, GenePT, DepMap). Each point is one perturbation, colored by its train/validation/test assignment. The first four columns are single-gene perturbations, each represented by its own embedding vector; the Norman column contains double perturbations, each represented by the mean of its two genes’ embedding vectors, with marker shape denoting the GEARS evaluation subgroup (unseen single, and combinations with zero/one/two genes seen in training). Across all datasets and embedding methods, the train, validation, and test perturbations are thoroughly intermixed rather than occupying distinct regions. However, there are clearly separated tiny clusters in the Norman datasets in three embedding spaces.

